# Long-lived IgE plasma cells that reside in the spleen contribute to the persistence of the IgE response

**DOI:** 10.1101/2025.09.18.676877

**Authors:** Mariana C.G. Miranda-Waldetario, Edgar Gonzalez-Kozlova, Kenneth B. Hoehn, Edenil Costa Aguilar, Laura Xie, Carlos J. Aranda, Yolanda Garcia-Carmona, Erica G. M. Ma, Maria A. Curotto de Lafaille

**Affiliations:** Precision Immunology Institute, Icahn School of Medicine at Mount Sinai (ISMMS), New York, NY 10029; Jaffe Food Allergy Institute, ISMMS, New York, NY 10029; Department of Immunology and Immunotherapy, ISMMS, New York, NY 10029; Department of Biomedical Data Science, Geisel School of Medicine at Dartmouth, Hanover, NH 03755, USA; Department of Computational Biology, Weill Cornell Medicine New York, NY 10065, USA; MINDICH Child Health and Development Institute, ISMMS, New York, NY 10029; Department of Biochemistry & Molecular Pharmacology, New York University School of Medicine, New York, NY, 10016; Department of Pediatrics, ISMMS, New York, NY 10029

**Author notes:** Correspondence (M.A.C.L.) (M.C.G.M.W).

**Keywords:** IgE plasma cells, long-lived plasma cells, allergy, timestamping

## Abstract

IgE plasma cells (PC) producing high affinity antibodies are critical players in allergic diseases. Allergies may persist or resolve over time, but the factors involved in their evolution are not well known. Sustained production of IgE antibodies, even in the absence of allergen exposure in persistent allergy, suggests the existence of long-lived IgE PC. However, the ability of IgE PC to undergo terminal differentiation and become long-lived has been questioned. Here we demonstrate that IgE PC undergo swift maturation into non-dividing MHCII^low^CD93^+^CD98^high^ PC in immunized mice. Mature IgE PC have a distinct transcriptional profile for adaptation to high protein synthesis, glycosylation, and survival. Using PC timestamping, long-lived IgE PC could be found several months after immunization in mice. Remarkably, the spleen rather than the bone marrow, was a main tissue of residence of mature long-lived IgE PC. Our findings provide key insights to understand IgE production in persistent allergy.

**Highlights:**

- IgE PC form in a narrow window after immunization and undergo swift maturation.
- Timestamping reveals the existence of CD93^+^CD98^high^MHCII^low^ long-lived IgE PC.
- The spleen and lymph nodes are the main tissues of mature IgE PC residence.
- IgE PC use distinct adaptations to high antibody secretion and survival.

## INTRODUCTION

High affinity IgE antibodies mediate allergic diseases due to their ability to bind to high-affinity FcεRI receptors on mast cells and to induce mast cell degranulation upon cross-linking by allergens.^1^ The production of IgE antibodies is highly restricted in several ways.^2–4^ IgE producing cells exist mainly as plasma cells (PC),^5^ as the IgE germinal center phase that develops in primary responses is short lived and IgE memory B cells are scarce or non-existent.^6,7^ The production of high-affinity IgE PC relies mostly on IL4R-dependent sequential switching to IgE of affinity matured type 2 IgG1 memory B cells.^6,8–13^

IgE PC have, for long, been considered immature and short-lived. They were found to be unfit to migrate to the bone marrow (BM), rather remaining in lymphoid organs such as lymph nodes (LN) and spleen.^5,7^ Poor localization of IgE PC in the BM was attributed to their impaired migratory response to CXCL12.^14^ Human IgE PC from blood^15,16^ and from the nasal mucosa^17^ were described as immature based on the expression of MHCII and CD23. In a mouse model of food allergy, IgE antibodies able to mediate anaphylaxis were greatly decreased after a primary immunization, supporting the notion that IgE PC are short lived.^18^ The seasonal variations of IgE antibodies to aeroallergens^19^ also argues for short life of IgE PC. Furthermore, inhibition of IL4R signaling in humans and mice, which prevents the formation of new IgE PC, led to a large decrease in circulating IgE^20–22^ and in decrease anaphylactic responses.^23^

Membrane IgE, the IgE B cell receptor (BCR), is a main driver of IgE cell fate. Forced expression of membrane IgE induces PC differentiation^24,25^ and promotes apoptosis through the Syk-BLNK-JNK/p38 pathway.^24^ Chronic Ca2+ signaling by the IgE BCR contributes to cell death.^26^ The cytoplasmic tail of membrane IgE was found to promote apoptosis by binding to Hax1.^27^ Furthermore, crosslinking of the IgE BCR by anti-IgE antibodies or antigens was described to lead to IgE PC death.^28^

However, in human atopic individuals treated with blocking IL4R antibodies, circulating IgE remains at about 40% levels of pre-treatment, which suggests that part of the IgE PC population is long lived.^20–22^ Other observations also support the existence of long-lived IgE PC. For example, IgE against foods persists in many food allergic individuals that do not consume the allergenic food.^29^ Individuals that were infected with the parasite filaria in an endemic area, moved to a non-endemic country and were treated with anti-helminth therapy, continued to have IgE antibodies against the parasite after several years.^30^ An early study in mice reported persistence of IgE and IgG responses up to one year after immunization, even after the depletion of memory B cells by X-irradiation.^31^ In another study of mice immunized intraperitoneally and challenged repeatedly through the airways, IgE PC were found in spleen and BM after 100 days of the last antigen administration.^32^ Repeated immunizations with house dust mite extract led to the formation of long-lived BM IgE PC that were detected up to a year after the last antigenic challenge.^33^ Multiple antigen exposures are in general necessary for the development of high affinity IgE antibodies^5^ and may also be required for the development of long-lived IgE PC.

Here we investigated the maturation, localization and life span of IgE PC in mouse models of type 2 responses. We found that IgE PC formed as plasmablasts swiftly undergo maturation into MHCII^low^CD93^+^ non-dividing PC. Most new and mature PC were found in LN and spleen and few mature IgE PC resided in the BM. This contrasted with IgG1 PC, which formed over a longer period of time, and readily colonize the BM. Using timestamping of PC, we demonstrated the existence of long-lived IgE PC with a mature phenotype. IgE PC transcriptional profile indicated a robust adaptation to the cellular stress originated by high antibody synthesis and N-glycosylation. In sum, this study demonstrates that IgE PC undergo maturation, and some become long-lived, and that mature long-lived IgE PC reside mainly in secondary lymphoid organs rather than in the BM. These findings offer new insights into the persistence of allergic responses.

## RESULTS

### IgE PC undergo maturation in secondary lymphoid organs and bone marrow

To characterize IgE PC *in vivo*, we used TBmc mice, which have monospecific populations of CD4 T and B cells that are specific for chicken ovalbumin (OVA) and for a linear peptide from influenza virus hemagglutinin (HA) respectively.^34^ TBmc mice are a good model to study IgE cells because they have augmented type 2 responses to immunization.^5,34^ T cells of TBmc mice are specific for a peptide from TBmc mice are typically immunized with OVA crosslinked to a mutated hemagglutinin peptide (PEP1), which does not bind to their naïve B cells.^5^ In this way, affinity maturation of the antibody response can be tracked by the appearance of PEP1-specific antibodies and PEP1 affinity–enhancing mutations in the V regions of immunoglobulin (Ig) genes.^5,8^

To characterize the transcriptome of PC induced by immunization in conditions favoring the accumulation of IgE PC of high affinity,^5^ TBmc mice were immunized 3 times with OVA-PEP1 in alum by intraperitoneal route. Three weeks after the last immunization, IgM-negative PC were isolated from bone marrow (BM), mesenteric lymph nodes (LN), and spleen (SP), and subjected to single cell RNA sequencing (scRNAseq) using the 5’ 10X Genomics platform for the analysis of whole transcriptome expression and B cell receptor (BCR) repertoire (Figure 1A). A total of 3,319 PCs with assembled BCRs were recovered across all tissues. The distribution of cells expressing IgGs, IgA and IgE BCRs in tissues is shown in Figure 1B. IgG1 PC were the most numerous, followed by IgE and IgA PC, and by lower numbers of IgG2a and IgG2b PC. Very few IgG3 PC were present in the dataset (not shown). IgG1 PC were comparably distributed among BM, LN and spleen, while IgA, IgG2a and IgG2c had similar distribution in spleen and BM but lower numbers in LN. IgE PC were distinctly distributed, being remarcably more numerous in spleen and LN than in BM (Figure 1B and Figure S1A).

**Figure 1.**
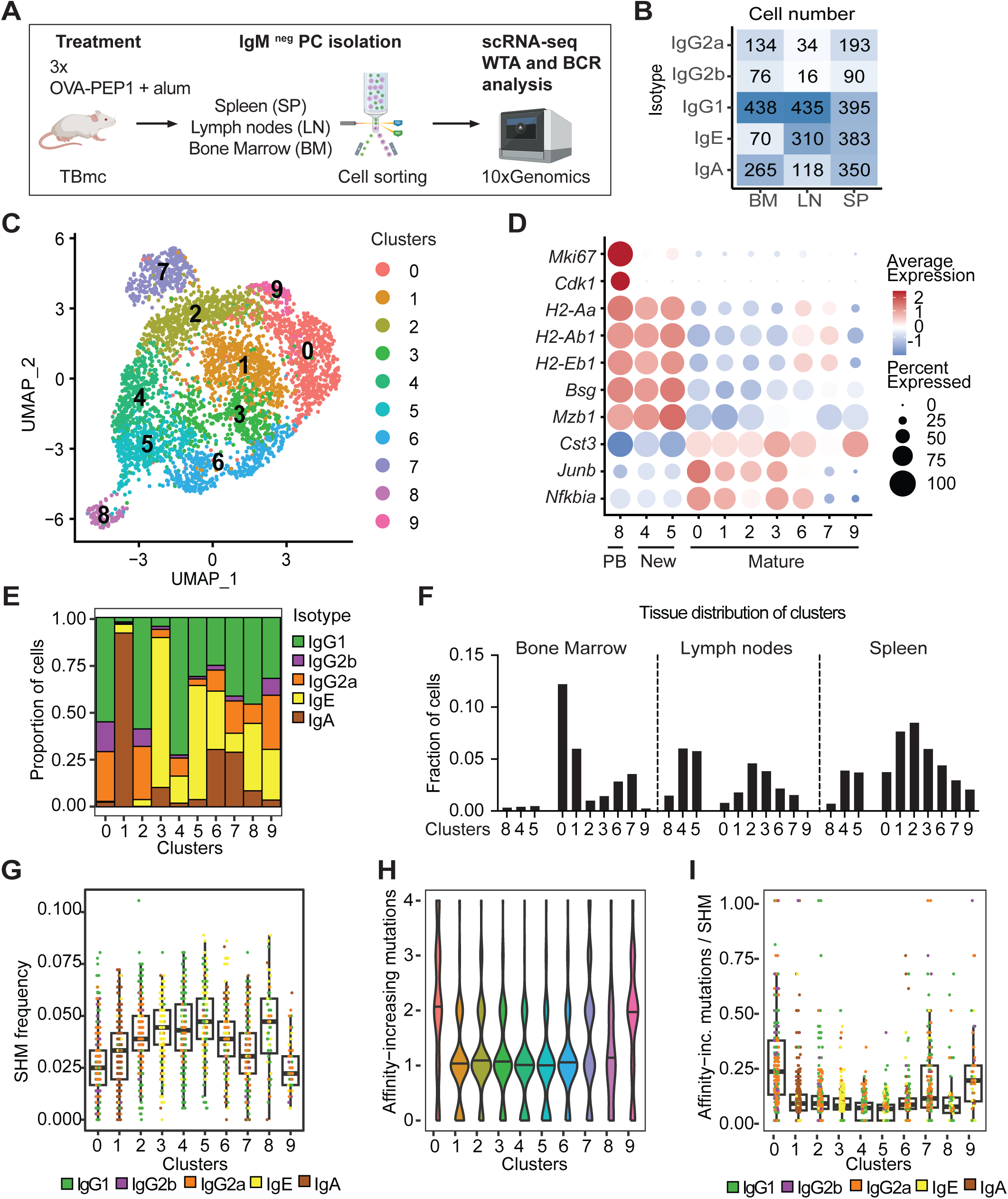
Transcriptional analysis of PC maturation (A) Workflow of single-cell RNA sequencing (scRNA-seq) analysis of PC from TBmc mice immunized three times with OVA-PEP1 in alum by intraperitoneal route. Three weeks after the last immunization, IgM^-^CD138^+^CD98^+^ PC were isolated from the spleen (SP), mesenteric lymph nodes (LN), and BM. The 5’ 10X Genomics platform was used for whole transcriptome and B cell receptor (BCR) sequencing. (B) Total number of sequenced cells per isotype and organ. (C) UMAP of PC clusters after Seurat analysis. (D) Dot plots of selected genes to identify clusters containing plasmablasts (PB), new PC and mature PC. (E) Proportion of PC expressing switched isotypes in each cluster. (F) Distribution of cells per cluster in bone marrow, lymph nodes and spleen. (G) Frequency of somatic hypermutations (SHM) per cluster, colored by isotype. (H) Violin plots of the distribution of BCRs carrying 0 to 4 high affinity mutations in their CDR3 heavy chain, per cluster. (I) Ratio of high affinity mutations to SHM frequency per cluster. See also Figure S1.

Using Seurat analytical tool with exclusion of immunoglobulin genes, we identified 10 distinct transcriptional clusters (Figure 1C). The PC identity of cells in the clusters was confirmed by high expression of typical PC genes *Prdm1*, *Xbp1*, *Jchain* and *Irf4*, and low expression of typical naïve/memory/germinal center B cell genes such as *Bach2* and *Bcl6* (Figure S1C-D).

To determine the differentiation status of cells in the clusters, we analyzed differentially expressed genes (Table S1). B cell differentiation into PC involves upregulation of Blimp-1, which in turn negatively regulates the expression of MHCII genes.^35^ New PC emerge as B220^hi^MHCII^hi^ cells that can further differentiate into mature B220^lo^MHCII^lo^ and became long-lived PC.^36^ Plasmablasts are recently formed, proliferating PC. To identify plasmablasts, new PC and mature PC, we analyzed the expression of cell division associated genes *Ki67* and *Cdk1* and MHC II genes *H2-Aa, H2-Ab1,* and *H2-Eb1* (Figure 1D). Cluster 8 had high expression of Ki-67 and other cell division genes as well as MHCII genes, thus contained plasmablasts (PB). Clusters 4 and 5, which contained non-dividing PC with high MHCII expression, were identified as recently formed ‘new’ PC. All other clusters, 0, 1, 2, 3, 6, 7, and 9, which contained non-dividing PC with lower average expression of MHC II genes, were classified as mature PC. Other differentially expressed genes of plasmablasts and new PC clusters included *Bsg* and *Mzb1*, while *Cst3* was differentially expressed in all mature clusters, and *Junb* and *Nfkbia* were differentially expressed in mature clusters 0, 1, 2, 3 and 6 (Figure 1D and Table S1).

We then analyzed the isotype distribution across clusters and tissues using BCR data (Figure 1E-F and Figure S1A-B). Some clusters had a dominant isotype composition. For example, cluster 0 and 2 contained mostly IgGs PC with predominance of IgG1 PC; cluster 1 had predominantly IgA PC; cluster 3 had predominantly IgE PC; and cluster 4 had predominantly IgG1 PC (Figure 1E). Other clusters had mixed isotype composition (Figure 1E). BM PC where mostly found in clusters of mature PC such as clusters 0,1, 6 and 7, whereas PC from spleen were found in all clusters, and LN PC had higher frequency in clusters 2, 3, 4, 5 and 8 (Figure 1F). IgE PC were present in early clusters 8, 4 and 5 and the mature clusters 3 and 6, and few IgE PC were found in clusters 7 and 9. BM IgG1 PC were predominantly found in cluster 0, followed by cluster 7. BM IgA PCs were mostly found in clusters 1 and 7 (Figure 1E and Figure S1A).

The analysis of the frequency of somatic hypermutations (SHM) and of high affinity mutations^5,8,10^ (Figure 1G-I and Figure S1E-G) reveled that plasmablasts (cluster 8) and new PC cluster 5 had the highest SHM frequency, and clusters 1 (IgA PC) and new PC clusters 4 and 5 had the lowest frequency of high affinity mutations (Figure 1G). On the other hand, mature clusters 0, 2, 7 and 9 had high frequency of high affinity mutations, which were highest in cluster clusters 0 (IgG PC of BM) (Figure 1H). In fact, the ratio of high affinity mutations to SHM was highest for mature clusters 0, 7 and 9 and lowest for new clusters 4 and 5 (Figure 1I). Thus, new PC appeared to be formed from highly mutated low affinity clones, while mature PC contain less mutated clone of higher affinity. This can be explained by the inhibitory effect of pre-existing high affinity antibodies on the activation of high affinity clones after a recall immunization.^37^

In sum, we identified new and mature PC in a mouse model of multiple immunizations. Maturation of the PC response was characterized by exit from the cell cycle and downregulation of MHCII genes. IgE PC underwent maturation, but mature IgE PC were preferentially found in secondary lymphoid organs such as the spleen and LN.

### IgE PC have increased expression of genes of ER stress, protein synthesis and glycosylation

To identify transcriptional differences between IgE and IgG1 PC, we grouped the PC of each of these isotypes from new or mature PC clusters. Plasmablasts were excluded as their transcriptional profile is dominated by cell division gene expression. Since spleen and LN PC of each isotype were found in overlapping regions of the UMAP (Figure S1B), spleen and LN (spleen+LN) PC of each isotype were grouped. The projections of new and mature IgE, IgG1 and IgA PC in the UMAPs of spleen+LN and BM are shown in Figure 2A. Spleen+LN contained new and mature IgE and IgG1 PC while BM PC of the two isotypes were mostly mature PC (Figure 2A). We then compared the transcriptional profile of IgE and IgG1 PC from the groups of: new PC from spleen+LN, mature PC from spleen+LN, and mature PC from BM (Figure 2B and Table S2). Overall, differences between IgE and IgG1 PC involved higher expression of many differentially expressed genes (DEGs) in IgE PC compared with IgG1 PC. Among those, we observed higher expression of *Slc3a2*, encoding for the amino acid transporter CD98, *Fcer2,* encoding for the IgE low affinity receptor CD23, and *IL-13R*α. In addition, new IgE PC displayed high expression of *Rgs1*, a regulator of G protein signaling and CXCL12-mediated migration.^38^ In both isotypes, there was a tendency towards transcriptional downregulation of genes along maturation in spleen+LN. BM IgE PC had a distinct transcriptional profile with notable upregulation of a group of genes exclusively in this population, and upregulation of another group of genes in mature IgE PC of both spleen+LN and BM (Figure 2B and Table S2). Genes highly expressed in IgE PC compared to IgG1 PC included those of the endoplasmic reticulum (ER) stress and unfolded protein response (UPR) (Figure 2C), protein synthesis, and N-glycosylation (Figure 2D). Nevertheless, these pathways tended to attenuate during maturation in both isotypes (Figure 2C-D). Consistently, using the ModuleScore function, we found that the ER stress score was highest in new IgE and IgG1 PC from spleen+LN cells, and lowest in mature cells of spleen+LN and BM (Figure 2E). The translation score was also higher in new versus mature IgE PC of spleen+LN and BM, and in new versus mature IgG1 PC of spleen+LN (Figure 2F). The N-glycosylation score was highest in IgE than IgG1 cells and decreased with maturation in both isotypes (Figure 2G). This adaptation of IgE PC may respond to the high N-glycosylation demand for IgE antibodies.^39^ Consistent with the increased translational score and ER stress response of IgE PC, we found that when cultured in the presence of APRIL and IL-6,^40^ IgE PC tended to secrete more antibodies on a per cell basis than IgG1 PC (Figure S2).

**Figure 2.**
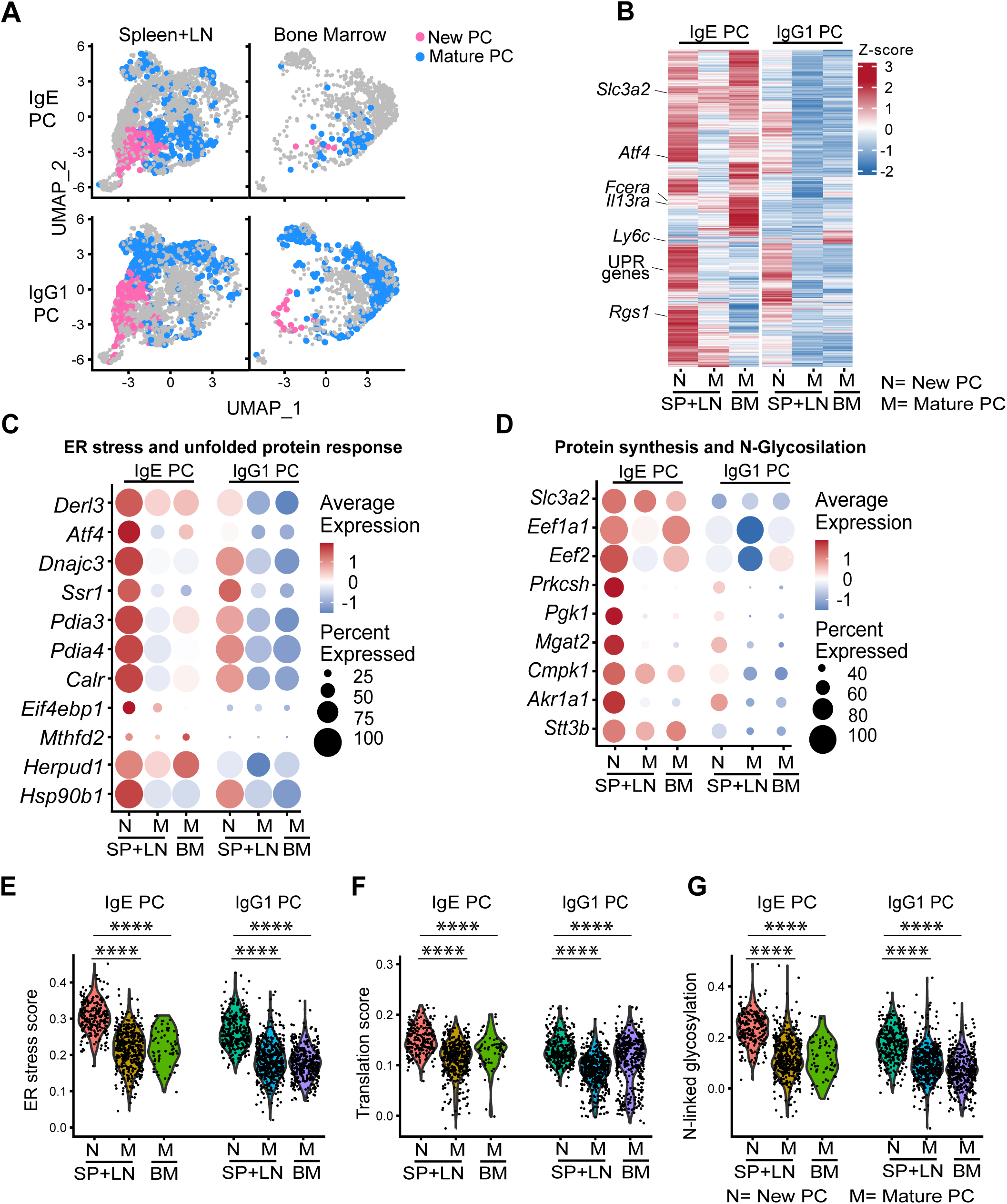
IgE PC transcriptional profile denotes distinct adaptation to the secretory function (A) Projection of new and mature IgE and IgG1 PC from spleen+LN and from BM on the UMAP of all PC. (B) Heatmap of differentially expressed genes (p ≤ 0.05) between new (N) and mature (M) IgE and IgG1 plasma cells (PC) from spleen and lymph nodes (SP+LN) and mature (M) IgE and IgG1 plasma cells from bone marrow (BM). UPR: unfolded protein response. (C) Dot plots of selected genes associated with the ER stress and unfolded protein response. (D) Dot plots of selected genes associated with protein synthesis and N-glycosylation. (E-G) Violin plots showing gene signature scores for ER stress (E), Translation score (F), N-linked glycosylation (G). Statistical analysis was performed using one-way ANOVA; ****: p≤ 0.0001. See also Figures S2 and S3.

In agreement with the attenuation of the ER response with maturation, pseudotime analysis of IgE and IgG1 PC (Figure S3A-C) revealed that high pseudotime score correlated negatively with the expression of ER stress genes (Figure S3B). Furthermore, in the clusters’ analysis, the overall ER stress score was highest in plasmablasts and new PC clusters (clusters 8, 4 and 5) than in mature PC clusters (Figure S3C). Overall, the results indicate that even though there is attenuation of cellular stress and protein synthesis pathways with maturation of IgE PC, these pathways remain more elevated in IgE PC than other PC.

### Mature IgE PC upregulate expression of survival genes

To assess the potential for long term persistence of mature IgE PC, we analyzed the expression of genes associated with PC survival and lifespan. It was recently shown that BM IgE PCs exhibit increased expression of pro-survival genes.^41^ We found that mature IgE PC of spleen+LN and of BM, compared with new IgE PC, expressed higher levels of survival genes *Tnfrsf17* (encoding BCMA), *Tnfrsf13b* (encoding TACI), and *JunB*, as well as a set of genes (*Pim1, Gpx4, and Lars2*) known to promote cell survival under cellular stress (Figure 3A).^42–44^ Compared to other PC, mature BM IgE PC expressed elevated levels of *Bcl2* and *Mcl1,* critical anti-apoptotic genes.^45^ *CD93*, encoding C1qR1, a protein required for maintenance of antibody secretion in PC,^46^ was highly expressed in about 20% of BM IgE PC. When mature IgE PC were segregated by clusters, IgE PC from cluster 6 had the highest expression of survival genes (Figure S4). Projection of IgE PC expressing *Bcl2* and *Mcl1* in the UMAP demonstrated the localization of *Bcl2*+ IgE PC in mature clusters 6 and 3 (Figure 3B), while Mcl1+ IgE PC were more broadly distributed (Figure 3C). Expression of some survival genes (e.g., *Tnfrsf17* and *Tnfrsf13b*) was higher in mature IgE PC than mature IgG1 PC, which may indicate that different PC isotypes rely on different survival strategies.

**Figure 3.**
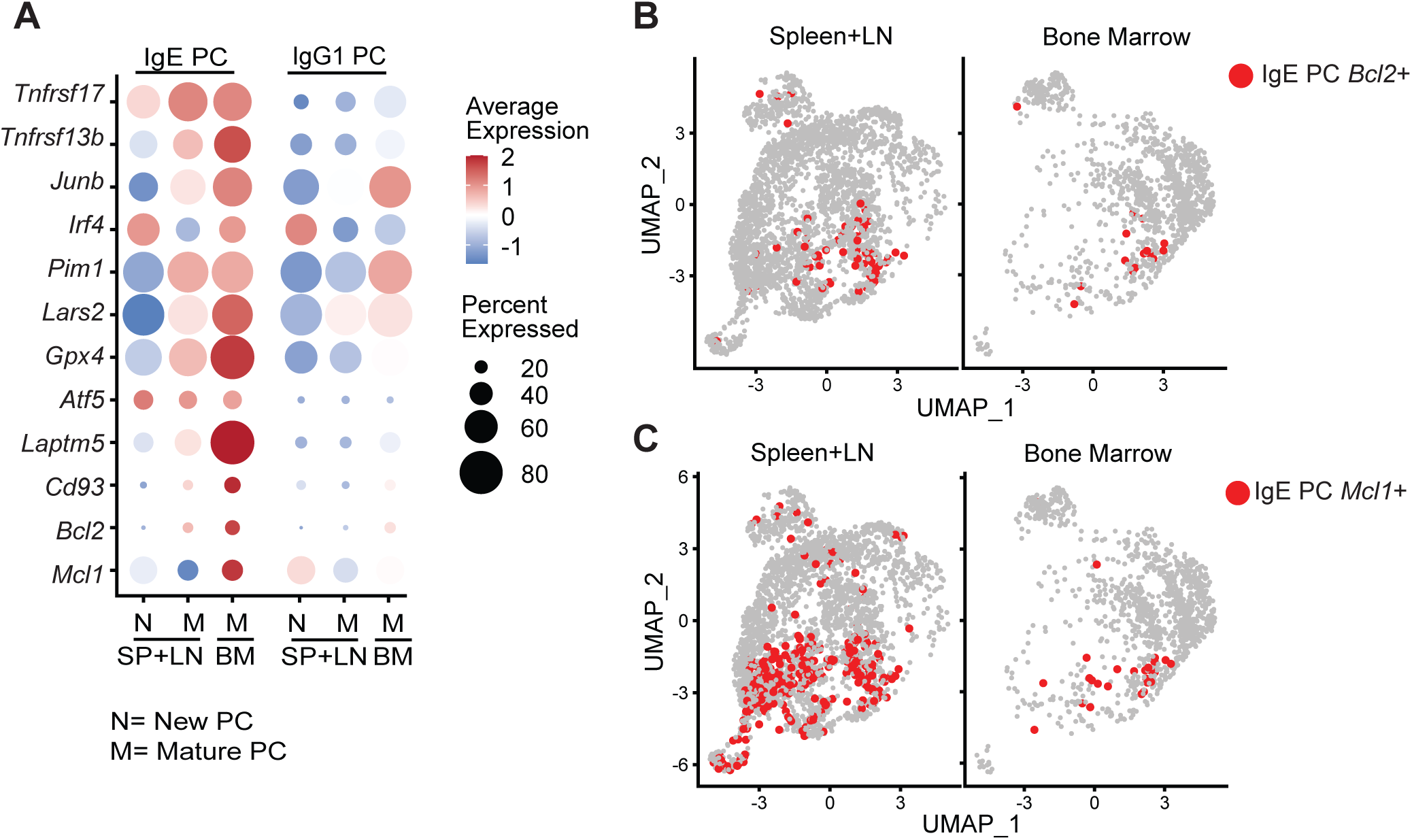
Cell survival signatures of mature IgE PC (A) Dot plot graphics of selected genes that were previously associated to PC survival. (B) Projections of Bcl2+ IgE PC from spleen+LN (left) and from BM (right) on the UMAP of all PC. (C) Projections of Mcl1 + IgE PC from spleen+LN (left) and from BM (right) on the UMAP of all PC. See also Figure S4.

We previously described that circulating IgE PC isolated from asthma and atopic dermatitis (AD) patients differentially expressed *Atf5*,^16^ an ER stress response genes associated with secretory cells’ survival,^47^ and *Laptm5*, a negative regulator of the BCR expression and B cell activation.^48^ *Laptm5* was highly expressed by mature BM IgE PC (Figure 3A). In sum, we demonstrated that the maturation of IgE PC involves upregulation of genes important for PC survival.

### IgE PC form early after primary immunization and accumulate in the spleen

To better understand the kinetics of IgE PC formation and persistence, we analyzed PC from spleen and BM of TBmc mice for over 150 days after primary immunization (Figure 4A). Total IgG1 and IgE levels increased after immunization and remain elevated by day 150 after immunization (Figure 4B). As previously described,^5^ circulating PEP1-specific IgE antibodies were detected later than PEP1-specific IgG1 antibodies. Furthermore, PEP1-specific IgE antibody titers at peak on day 70 were about 10-fold lower than PEP1-specific IgG1 antibody titers (Figure 4C).

**Figure 4.**
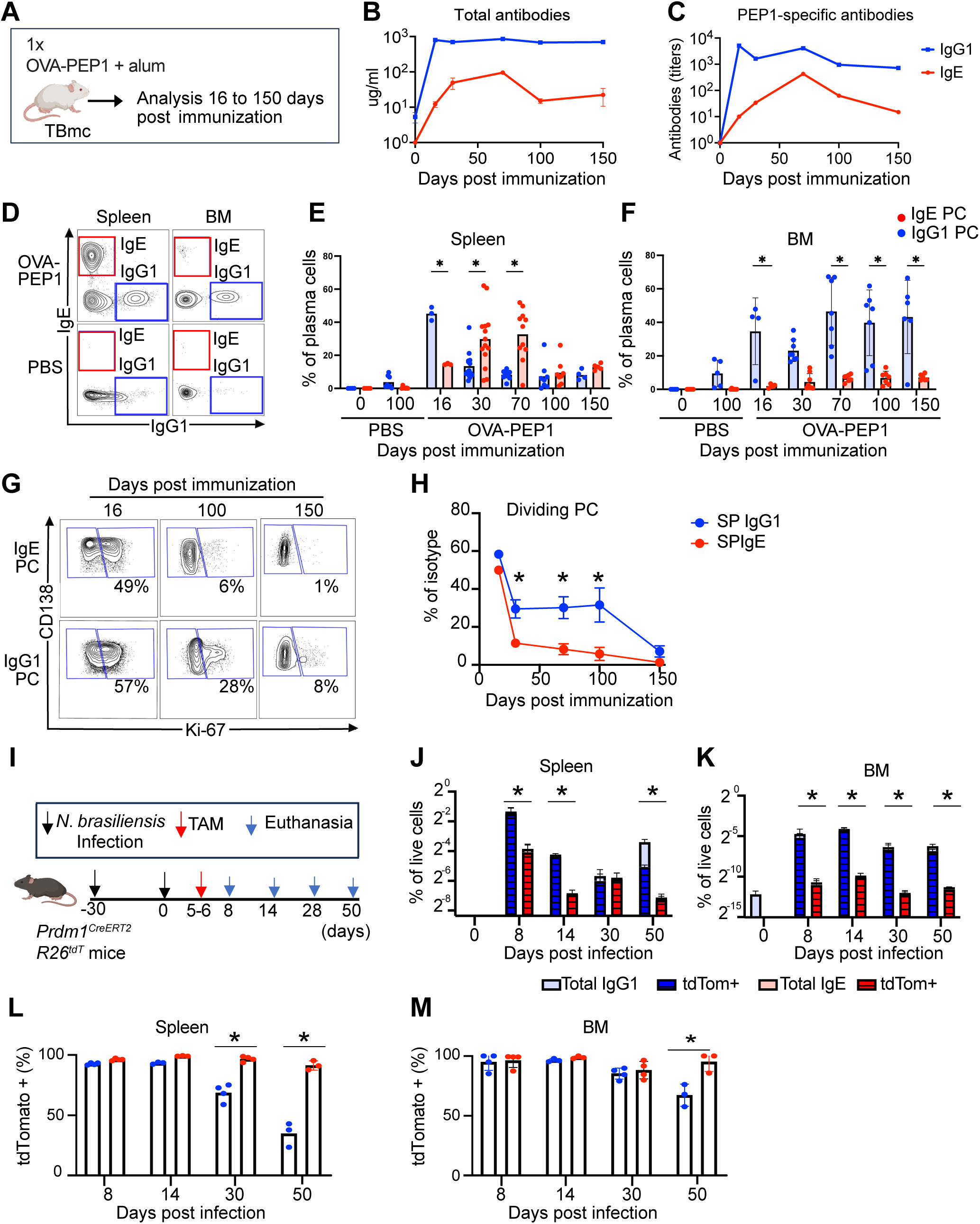
Kinetics of formation and maturation of IgE PC (A) Experimental design for data in (B to H). TBmc mice were immunized once by intraperitoneal route with OVA-PEP1 in alum, and PC and antibodies were analyzed from days 16 to 150 after immunization. (B) Concentration of total IgG1 and total IgE antibodies in plasma (n=3-4 per timepoint). (C) Titers of PEP1-specific IgG1 and IgE antibodies in plasma (n=3-4 per timepoint). (D) Representative flow cytometry contour plots illustrating detection of IgE PC and IgG1 PC among PC (gated as CD138^+^CD98^+^) from OVA-PEP1 immunized mice (day 30 post-immunization) and unimmunized PBS-injected mice. (E-F) IgE PC and IgG1 PC frequencies among spleen PC (E) and BM PC (F). Data is from three independent experiments. Each dot corresponds to one mouse sample. (G) Ki-67 staining of IgE PC and IgG1 PC at 16, 100, and 150 days post-immunization. (H) Percentage of Ki-67^+^ cells among IgE PC and IgG1 PC at the indicated time points. Data is from two combined experiments (n = 3–5 mice per time point). (I) Experimental design for data in (J to M). *Prdm1^CreERT2^R26^tdT^* mice were infected subcutaneously with *Nippostrongylus brasiliensis* on days -30 and 0, and administered tamoxifen (TAM) on days 5 and 6 to timestamp PC. Euthanasia was performed on days 8, 14, 28, and 50 post-secondary infection. (J-M) Percentage of tdTomato^+^ among total IgG1 PC (blue) and IgE PC (red). In the spleen (J) and BM (M). n = 3–4 mice per time point. Mean ± SEM are shown. Statistically significant differences, determined using unpaired Student’s *t*-test, are indicated (*:p ≤0.05). See also Figure S5 and S6.

IgE and IgG1 PC in spleen and BM were analyzed by flow cytometry (Figure S5A and Figure 4D). Total PC in the spleen reached their highest numbers by days 16-30 and decreased thereafter (Figure S5B). As expected, PC frequency increased later in the BM, peaking by day 100 after immunization (Figure S5C), consistent with the formation of PC in secondary lymphoid organs and subsequent migration to the BM.^49^ PC frequency was very low in unimmunized TBmc mice but increased over time in spleen and BM (days 0 and 100, PBS) (Figure S5B-C).

We next determined the frequency of spleen and BM IgE and IgG1 PC along the 150 days post immunization (Figure 4D-F). IgE and IgG1 PC were undetectable in spleen and BM of most unimmunized mice on day 0 (Figure 4E-F and Figure S5D-E). IgG1 PC frequency was highest at day 16 of immunization in spleen and decreased thereafter (Figure 4E). In BM, IgG1 PC increased by day 16 of immunization and remained at relatively constant levels thereafter (Figure 4F).

Consistent with the IgE antibody response, IgE PC peaked between days 30-70 in spleen, when they constituted close to 40% of all PC, greatly surpassing IgG1 PC numbers. Spleen IgE PC declined by days 100-150 post immunization (Figure 4E). There was delayed and very low accumulation of IgE PC in the BM over the 150 days post immunization (Figure 4F). To determine if new formation of IgE PC could account for their accumulation in the spleen, we analyzed the kinetics of Ki-67 expression after immunization (Figure 4G-H). The highest percentages of Ki-67^+^ IgG1 and IgE PC were observed at day 16 of immunization, when intense PC formation was occurring, and at least half of both PC isotypes expressed Ki-67 (Figure 4G-H). Ki-67^+^ IgE PC decreased drastically by day 30 of immunization to almost undetectable levels by day 150, indicating that most IgE PC were formed in the first 30 days after immunization. In contrast, 30% or more of IgG1 PC in spleen were Ki-67^+^ up to day 100 post immunization, suggesting a continuous generation of IgG1 PC over a prolonged period of time. The continuous production of IgG1 PC without accumulation in the spleen, together with their sustained numbers in the BM, suggest that a swift migration of new IgG1 PC to the BM maintained the IgG1 BM population.

To further investigate the kinetics of IgE and IgG1 PC formation, we used the *Prdm1^CreERT2^R26^tdT^* mouse strain generated in our laboratory (Figure S6A), which allow to timestamp nearly all PC at a chosen time by administration of tamoxifen (Figure S6B-C). Following tamoxifen administration, virtually all already formed PC will constitutively express dtTomato, while new PC formed after tamoxifen is cleared are dtTomato negative. This model also allows to analyze PC differentiation responses in mice with polyclonal T and B lymphocyte repertoires.

*Prdm1^CreERT2^R26^tdT^* mice were infected with the parasite *Nippostrongylus brasiliensis,* which induces transient IgG1 and IgE PC responses.^6^ *Prdm1^CreERT2^R26^tdT^*mice were in the C57Bl/6 genetic background which is less prone to type 2 and IgE responses.^50^ To increase the number of PC we analyzed PC after a secondary infection. Tamoxifen was administered on days 5-6 post re-infection, and PC were analyzed on days 8, 14, 30 and 50 thereafter (Figure 4I). SP IgE PC were less abundant than SP IgG1 PC at all time points analyzed, except at day 30 post immunization (Figure 4J). Very few IgE PC were observed in the BM after infection (Figure 4K). At day 8 post infection, more than 95% of all SP and BM IgG1 and IgE PC were tdTomato+(Figure 4L-M). Interestingly, the percentage of tdTomato+ IgG1 PC in spleen declined to about 70% on day 30, and to less than 40% on day 50, reflecting the formation of new, tdTomato negative IgG1 PC (Figure 4L and Figure S6D). In contrast, IgE PC that remained up to day 50 post infection were more than 90 % tdTomato+. IgG1 turnover was also observed in the BM but to a lesser extent than in the spleen, with about 60% of the IgG1 BM being tdTomato+ 50 days after infection (Figure 4M and Figure S6E).

Together, these results indicate that IgE PC are produced mostly in a single early wave after immunization or infection, subsequently decreasing in numbers. Long-lived IgE PC persist mainly in secondary lymphoid organs and in very low numbers in the BM. In contrast, the formation of IgG1 PC is maintained for a much longer period during an immune response, and mature long-lived IgG1 PC are found less in secondary lymphoid organs and mostly in the BM.

### Mature MHCII^lo^CD93^+^ IgE PC are detected long after immunization in the spleen

To characterize the kinetics of appearance of early, mature, and long-lived spleen IgE PC after primary immunization, we analyzed spleen PC at various time points after immunization using spectral cytometer. Cells were stained with antibodies for PC identification and gating (CD138 and CD98), to determine PC maturation state (Ki-67 and MHCII), to determine isotype expression (IgM, IgA, IgG1, IgE), and expression of other described PC proteins,^51,52^ or proteins identified in our scRNAseq analysis (Table S1). tSNE-CUDA dimensionality reduction algorithms were used to identify populations within the CD98^+^CD138^+^ PC gate at 16 and 100 days after primary immunization (Figure S7A). Isotype-specific PC (IgM^+^, IgA^+^, IgG1^+^, and IgE^+^) were identified through manual gating and projected in the UMAP clusters of samples from day 16 and 100 after immunization (Figure 5A). The UMAP projections revealed little overlap between day 16 and day 100 PC, indicating occurrence of phenotypic shifts over time (Figure 5A). Each immunoglobulin isotype occupied a defined area of identity in the combined 16+100 days tSNE map (Figure 5B and Figure S7B). Ki-67^+^MHCII^high^ cells were characterized as PB, and Ki-67^-^ MHCII^high^ cells were characterized as early PC. PB and early PC were found among IgM^+^, IgG1^+^, and IgE^+^ PC.

**Figure 5.**
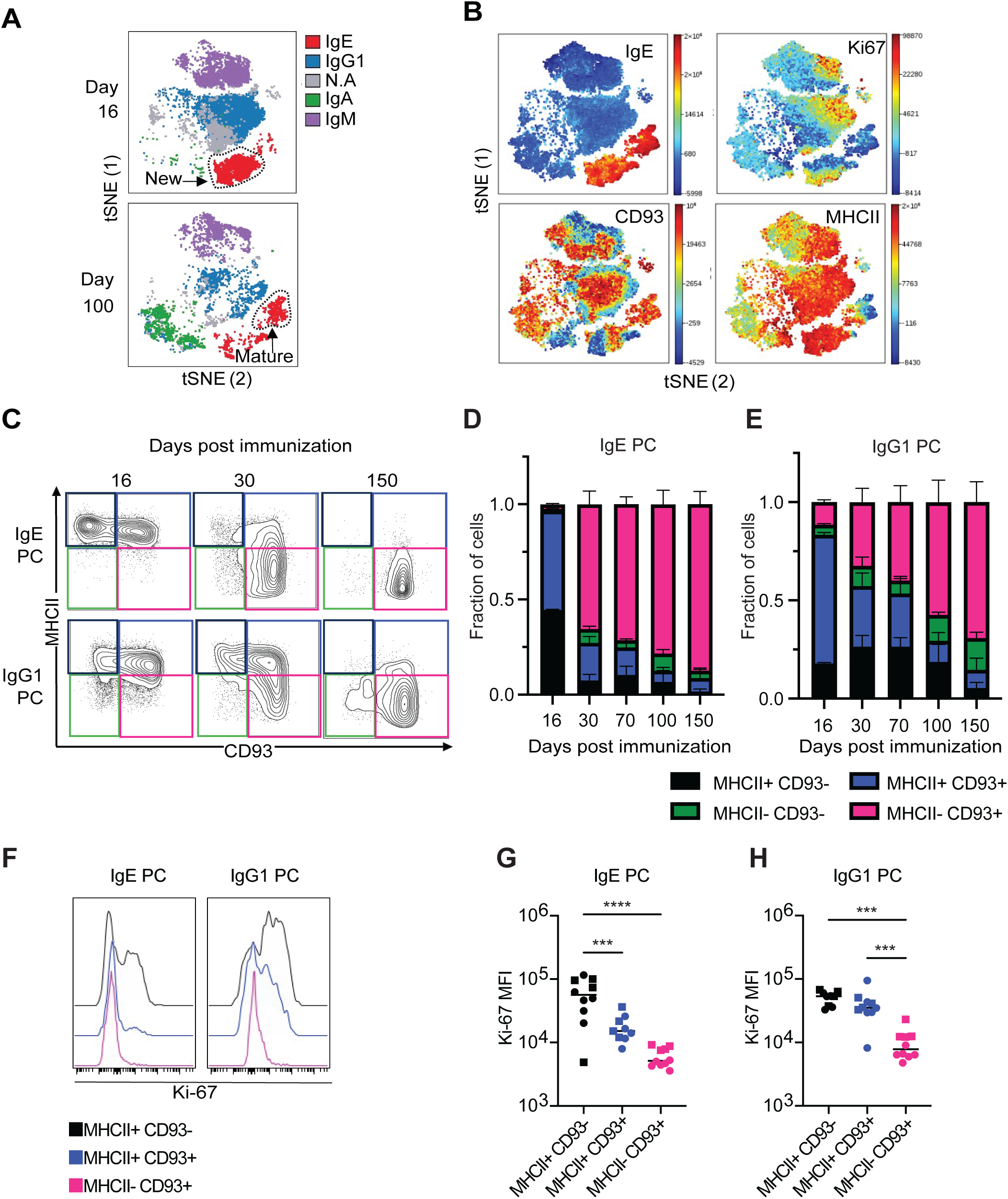
Mature IgE PC are MHCII^lo^CD93^+^ CD98^+^ (A) Projection of PC isotypes on concatenated tSNE plots of spleen PC from TBmc mice at 16 and 100 days post-immunization. Data is from spectral cytometry analysis. (B) Projection of IgE+, Ki67+, CD93+ and MHCII+ PC on tSNE plots of concatenated PC samples from spleen of mice at 16 and 100 days post immunization. (C) Expression of MCHII and CD93 in splenic IgE and IgG1 PC at 16, 30, and 150 days post-immunization defines 4 subpopulations: MHCII^+^CD93^-^, MHCII^+^CD93^+^, MHCII^-^CD93^-^and MHCII^-^ CD93^+^. (D-E) Changes in subpopulations of IgE PC (D) IgG1 PC (E) as defined by MHCII and CD93 expression, over 150 days post-immunization. Data are mean ± SEM from two combined experiments. n= 3–6 mice per timepoint. (F) Ki-67 expression in MHCII^+^CD93^-^, MHCII^+^CD93^+^, and MHCII^-^CD93^+^ subpopulations of splenic IgE and IgG1 PC. Data are from concatenated samples from days 30 and 150 after immunization. (G, H) Ki-67 mean fluorescence values in MHCII^+^CD93^-^, MHCII^+^CD93^+^, and MHCII^-^CD93^+^ subpopulations of spleen IgE PC (G) and IgG1 PC (H) from days 30 (squares) and 150 (circles) after immunization. Each dot represents one mouse. Statistical analysis was performed using one-way ANOVA; <u>p<</u> 0.01 is indicated as ***, and p≤ 0.0001 as ****. See also Figure S7.

Two areas of IgE PC were clearly distinguished in the tSNE plots, one corresponding to PB and early PC, which predominated at day 16, and the other corresponding to mature PC, which predominated at day 100 (Figure 5A-B). CD93 was consistently expressed in MHCII^low^ PC of all isotypes, indicating its association with the maturation of PC. However, CD93 was also expressed in some early PC, therefore additional markers, such as MHCII, were necessary to refine the identification of mature PC.

To assess differences in formation and maturation between IgE and IgG1 PC, we evaluated the changes in expression of MHCII and CD93 in splenic PC over 150 days following immunization. The combination of these two markers identified 3 distinct plasma cell populations MHCII^+^CD93^+^, MHCII^+^CD93^-^ and MHCII^-^CD93^+^ (Figure 5C). At 16 days post-immunization most IgE and IgG1 PC expressed MHCII, as expected for early PC,^36^ and part of the MHCII^+^ PC populations expressed CD93 (Figure 5C-E). Overtime MHCII^-^CD93^+^ emerged, presumably by downregulation of MHCII in MHCII^+^CD93^+^ PC. By day 150, most IgE PC were MHCII^-^CD93^+^, while a sizable population of MHCII^+^CD93^-^ IgG1 PC and some MHCII^+^CD93^-^ IgG1 PC were still observed. Ki-67 expression was highest in MHCII^+^CD93^-^ PC, followed by MHCII^+^CD93^+^ and was mostly absent in MHCII^-^CD93^+^ PC, confirming the latter mature status (Figure 5F). A large part of MHCII^+^CD93^-^ and MHCII^+^CD93^+^ IgG1 PC cells were dividing cells (Ki-67^+^) (Figure 5H), while less IgE PC were Ki-67^+^ in these populations (Figure 5G).

Other differences were observed between IgE and IgG1 PC (S7D-J). Unlike IgG1 PC, mature IgE PC did not upregulate EpCAM or Ly6C (Figure S7E-F). IgE PC on the other hand, expressed higher levels of CD98 than IgG1 PC, and part of the IgE PC population, but not the IgG1 PC population, express CD23 (Figure S7G-H).

This analysis demonstrated that IgE PC exit cell division and acquire a mature phenotype with a faster kinetics than IgG1 PC, consistent with the formation of IgE cells in an initial wave, in contrast to a prolonged generation of IgG1 PC, as evidenced by the sizable population of MHCII^+^Ki-67^+^ IgG1 PC even after 70 days post immunization (Figure 5E). These results validate the findings of the transcriptional analysis which indicated that IgE PC downregulate MHCII expression as they mature. Furthermore, the results demonstrate the differentiation of IgE PC into long-lived PC found up to 5 months after immunization.

We and others previously described that IgE PC formed at the peak of a primary response expressed higher levels of membrane IgE than IgE germinal center cells, contrary to the downregulation of membrane IgG1 in IgG1 PC.^6,7,53^ To determine if downregulation of membrane IgE occurs during PC maturation, we conducted an analysis using the Cytek Amnis ImageStream flow cytometer. IgE PC from the spleen of mice immunized 30 and 150 days earlier were surfaced stained with antibodies to CD138, MHCII, and Alexa 647-labeled anti-IgE (IgE SF). The cells were then permeabilized and intracellularly stained with Alexa488-labeled anti-IgE (IgE IC). IgE PC were gated by expression of CD138 and IgE IC. Varying levels of IgE SF and MHCII staining were observed among IgE PC (Figure 6A), including IgE PC with undetectable staining of IgE SF among the MHCII-low/negative cells, and less frequently among MHCII^+^ cells. Quantification of IgE SF on individual cells revealed that MHCII^+^ IgE PC expressed significantly higher levels of membrane IgE (Figure 6B). Cell size and IgE IC staining were comparable between MHCII^+^ and MHCII-low/negative cells (Figure 6C-D). These findings indicate that IgE PC downregulate membrane IgE expression during maturation.

**Figure 6.**
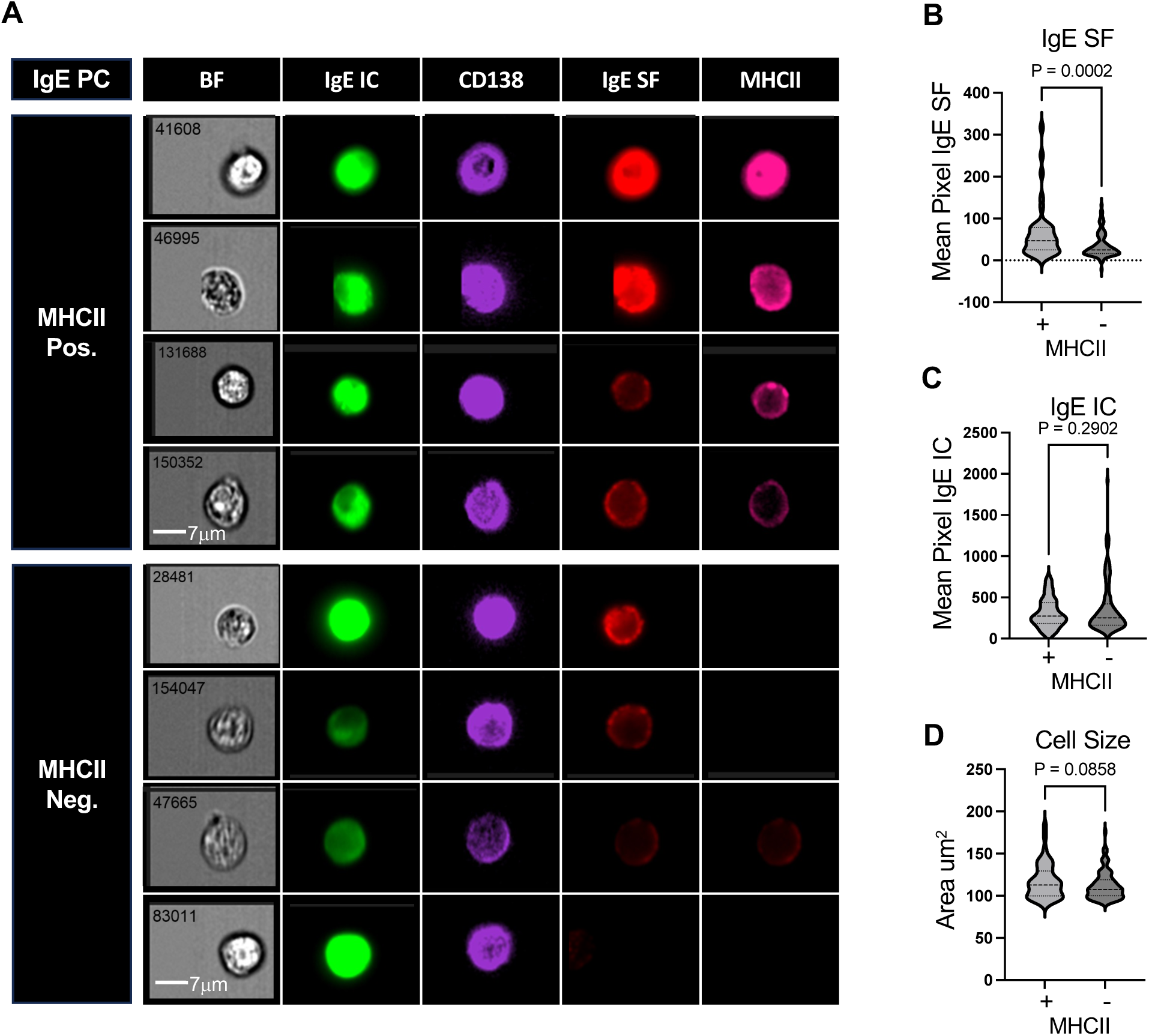
Mature IgE PC downregulate membrane IgE (A) Representative images from imaging flow cytometry (AMNIS) showing IgE PC, identified by co-expression of CD138 (purple) and intracellular IgE (IC, green), with varying levels of surface IgE (SF, red) and MHCII (pink). (B-D) Violin plots of 150 individual cell images gated as MHCII^+^ (+) or MHCII^low^ ^to^ ^negative^ (-). Violin plots of cell size measured by area (B). Violin plots indicating the average number of pixels corresponding to intracellular IgE (C). Violin plots indicating the average number of pixels corresponding to surface IgE (D). Statistical analysis was performed using an unpaired Student’s *t*-test; p-values are indicated.

### Long-lived IgE PC have a mature phenotype

To more accurately assess the persistence of long-lived IgE PC and to determine their phenotype, we used a model of allergic airway disease induced by the fungal extract *Alternaria alternata*,^54^ in combination with the timestamping *Prdm1^CreERT2^R26^tdT^*mouse model. *Alternaria alternata* extract was administered by intranasal route over a 10 week period to *Prdm1^CreERT2^R26^tdT^*mice. At the end of the immunization, we administered tamoxifen to label all PC present at that time point (Figure 7A). After 1 and 20 weeks, we evaluated the phenotype of PC in spleen and BM (Figure 7B-C and Figure S8). One week after tamoxifen administration 92% of BM PC and 95% of spleen PC were tdTomato+, confirming the efficiency of the PC timestamping method (Figure 7B). Over time, a higher turnover of PC was observed in the spleen compared to the BM: after 20 weeks, only 16% of spleen PC remained tdTomato+, whereas about 30% of BM PC remained tdTomato+ (Figure 7B) and. At week 20, the tdTomato negative fraction was a mix of PC aged between 1 day and 20 weeks and contained new and mature PC populations (MHCII^+^CD93^-^, MHCII^+^CD93^+^ and MHCII^-^CD93^+^). On the other hand, the 20 weeks old tdTomato+ fraction was formed by mature MHCII^-^CD93^+^ PC (Figure 7C). IgE antibodies in serum plateaued at about 8 weeks after the initiation of immunization and they decreased thereafter but still persisted at higher levels than those of pre-immunization at the 30 weeks timepoint (Figure 7D). Among tdTomato+ switched PC, IgE PC were more frequent in the spleen than the BM at 1 and 20 weeks after tamoxifen administration, consistent with our previous observations of maturation of IgE PC (Figure 7E).

**Figure 7.**
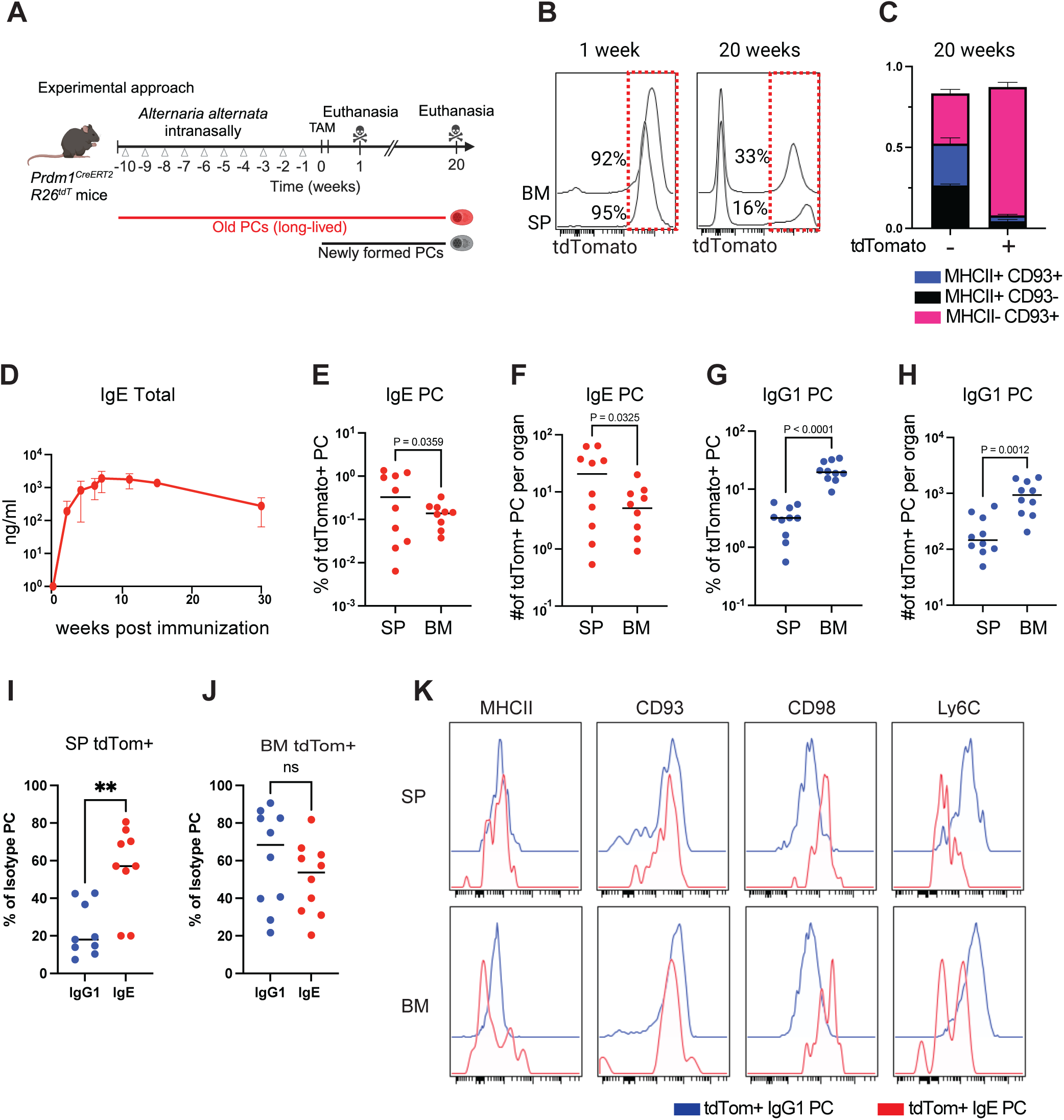
Timestamping reveals mature phenotype of long-lived IgE PC (A) Experimental design. *Prdm1^CreERT2^R26^tdT^* mice were chronically exposed to intranasal *Alternaria alternata* extract for 10 weeks and were then administered tamoxifen (TAM) every other day during the following week. Mice were euthanized at 1 and 20 weeks post-tamoxifen. (B) Average frequency of tdTomato+ PC in the spleen (SP) and bone marrow (BM) at 1 and 20 weeks post-tamoxifen. (C) MHCII^+^CD93^-^, MHCII^+^CD93^+^, and MHCII^-^CD93^+^ PC subpopulations among tdTomato negative and tdTomato+ PC at 20 weeks post-tamoxifen. Data are shown as mean ± SEM of 9 mice. (D) Plasma levels of total IgE. Data are shown as mean ± SEM of 9 mice. (E, H) Quantification of tdTomato+ IgE and IgG1 PC at 20 weeks post-tamoxifen. Percentages of IgE PC among tdTomato+ (tdT+) PC gate (E). Absolute numbers of tdTomato+ (tdT+) IgE PC per spleen (SP) and bone marrow from tibia-femur pair (BM) (F). Percentages of IgG1 PC among tdTomato+ (tdT+) PC gate (G). Absolute numbers of tdTomato+ (tdT+) IgG1 PC per spleen (SP) and bone marrow from tibia-femur pair (BM) (H). (I) Percentage of spleen (SP) tdTomato+ IgE and IgG1 PC among SP total IgE and total IgG1 PC, respectively. (J) Percentage of bone marrow (BM) tdTomato+ IgE and IgG1 PC among BM total IgE and total IgG1 PC, respectively. (E, J) Statistical analysis was performed using an unpaired Student’s *t*-test, with p-values indicated. (K) Expression of MHCII, CD93, CD98, and Ly6C in tdTomato+ IgE and IgG1 PC from spleen (SP) and bone marrow (BM). See also Figure S8.

Consitent with the preferential localization of mature PC in secondary lymphoid organs, tdTomato+ IgE PC were preferentially found in spleen than in the BM (Figure 7E). tdTomato+ IgG1 PC were as expected higher on the BM but were also observed in the spleen (Figure 7F). Consistent with our previous observation of continuous generation of IgG1 PC in the spleen, we found that at week 20 of tamoxifen administration, only about 14% of splenic IgG1 PC were tdTomato+, while about 60% of splenic IgE PC were tdTomato+. In contrast, the frequency of tdTomato+ PC was similar between IgE and IgG1 PC in the BM (Figure 7G-H).

Finally, we confirmed that long-lived tdTomato+ IgG1 and IgE PC in both the spleen and BM displayed a phenotype characterized as MHCII^low^CD93^+^. Furthermore, long-lived IgE PC expressed high levels of CD98 expression and low levels of Ly6C (Figure 7I) as described above for long-lived IgE PC in the TBmc model (Figure 5). These findings underscore the distinct characteristics and localization of long-lived IgE PC.

## DISCUSSION

IgE PC secrete the antibodies involved in triggering allergic reactions, including life-threatening anaphylaxis. The rarity of IgE PC has made their study challenging. Here we used experimental mouse models of type 2 immune responses, and employed scRNAseq, complex flow cytometry and in vivo timestamping to characterize the differentiation and lifespan of IgE PC. Contrary to the paradigm that IgE PC are immature and short-lived, we show that IgE PC reach maturation similarly to other PC. Furthermore, we demonstrate that long-lived IgE PC do develop, as they can be found several months after their formation, in agreement with previous findings.^31–33^ Strikingly, the spleen rather than the BM, is a major organ of residence of mature long-lived IgE PC.

PC are first formed as dividing plasmablasts in secondary lymphoid organs, and as they mature into specialized terminally differentiated secretory cells, they exit the cell cycle, downregulate typical B cell pathways such as those involved in antigen-presentation and BCR signaling, and upregulate the machinery necessary for antibody production and secretion. The ER stress, UPR, translation and glycosylation pathways are highly upregulated in PC. Mature PC are found in lymphoid organs and in the BM, where they survive in specialized niches. The BM is the organ of excellence for residence of long-lived PC.^36,55,56^ We demonstrate here that IgE PC that form in spleen and LN after immunizations undergo maturation, as evidenced by their exit from the cell cycle and downregulation of MHCII genes.^36,52^ We describe that maturation of PC involves downregulation of other genes, such as *Mzb1* and *Bsg*. Mature IgE PC upregulate genes common to mature PC of other isotypes, such as *Cst3*, *Junb*, *Nfkbia,* and genes involved in PC survival, such as *Bcl2*, *Mcl1*, *Tnfrsf17* (encoding BCMA),^57^ *Tnfrsf13b* (encoding TACI), *Pim1*,^58^ *Lars2*,^44^ and *Gpx4.*^43^ TACI and BCMA are receptors for BAFF and APRIL, important survival cytokines secreted by stroma cells in the BM,^59,60^ and by dendritic cells and macrophages in secondary lymphoid organs.^61^

As for other PC such as IgG1 PC, differentiation of IgE PC involved upregulation of molecular pathways necessary to adapt to the ER stress generated by high demand of protein translation, folding, N-glycosylation and secretion. We found that maturation of IgE and IgG1 PC, involves attenuation of cellular stress and protein synthesis molecular pathways. However, overall, expression of ER stress and translation pathways was augmented in IgE PC compared with IgG1 PC, as recently described.^41^ IgE antibodies are highly N-glycosylated compared with IgG1 antibodies,^39^ and glycosylation of IgE is important for pathogenesis.^62,63^ Consistently, N-glycosylation genes expression was higher in IgE PC than IgG1 PC. These results indicate that, while following general principle of PC maturation, IgE PC have distinct adaptations to antibody secretion and to survival.

IgE PC were found to differentially express *Atf5*, an ER stress response gene,^47^ *Laptm5*, encoding a transmembrane lysosomal protein controlling membrane BCR expression and B cell activation,^48^ *Rgs1*, an inhibitor of chemokine receptor signaling,^38^ *Fcer2*, encoding the low affinity IgE receptor CD23, and *IL13ra*. *Atf5*, *Laptm5* and *Fcer2* were also found to have higher expression in circulating human IgE PC than other PC.^16^

The significance of the expression of *Fcer2* and *IL13ra* in IgE PC is not yet known. It could however be a consequence of the ontogeny of these cells, as IgG memory B cells expressing *Fcer2* and other IL4R regulated genes have recently been found to be the memory precursors of IgE PC in persistent allergy.^11,12,16^ Increased expression of *Rgs1* in IgE PC may be in part responsible for their deficient response to CXCL12,^14^ the ligand for CXCR4, and for the poor establishment of IgE PC in the BM. CXCR4 has been shown to be necessary for the long-term homing of PC to the BM.^64^ Despite their deficient response to CXCL12 ^14^, the levels of CXCR4 in IgE PC are comparable to those of IgG1 PC.^41^

Differentiation of B cells into PC increases transcription of immunoglobulin genes and switches the ratio of membrane to secreted immunoglobulin mRNA. PC downregulate the production of membrane immunoglobulin while highly increasing the synthesis of the secreted form of immunoglobulin (antibody), a process regulated by differential mRNA polyadenylation and splicing.^65^ Previously, we and others determined that murine IgE PC formed at the peak of a primary response to immunization or helminth infection, expressed higher levels of membrane IgE than IgE germinal center cells, contrary to the decrease in membrane IgG1 observed in differentiating IgG1 PC.^6,7^ In this study we found that maturation of IgE PC involved downregulation of the IgE BCR, as MHCII^low^ IgE PC expressed significantly lower levels of membrane IgE than MHCII^+^ IgE PC. These differences may be due to transcriptional regulation of immunoglobulin isoforms, and/or to the increase in Laptm5 expression, which may promote lysosomal degradation of the IgE BCR.

We studied the kinetics of IgE PC formation, maturation, and survival. Recently formed PC were characterized as dividing - (Ki-67^+^MHCII^+^), new PC (Ki-67^-^MHCII^+^), and mature PC (Ki-67^-^ MHCII^low^). These three populations were identified among IgE and IgG1 PC of spleen and LN, while most BM PC had a mature phenotype. By analyzing PC over time using flow cytometry or combined flow cytometry and timestamping, we determined that IgE PC form in a relatively narrow period after immunization or infection, while there was a prolonged phase of IgG1 PC formation. Because of this, the IgE PC population tended to accumulate mature PC earlier than the IgG1 PC population, as the latter continued to contain PB and new PC for an extended time. Maturation of IgE, as well as other PC, involved upregulation of CD93,^52,66^ a receptor expressed in early B cell development and in PC, that is required for antibody secretion and PC persistence.^46^ Mature IgE PC could be distinguished from IgG1 PC by their higher expression of CD98, the amino acid transporter LAT1, and by low expression of Ly6C. Mature IgE PC can thus be characterized as MHCII^low^CD93^+^CD98^high^Ly6C^low^.

The lifespan of IgE PC has been a topic of active debate. IgE PC have been described as immature and short-lived, and unable to establish BM residence. After immunization or infection, only a small proportion of IgE PC reaches the BM, being mostly restricted to secondary lymphoid organs, while IgG1 PC readily colonizes the BM.^6,7^ In a model of food allergy, the half-life of the IgE PC population was calculated to be 60 days, much shorter than the 234 days half-life of IgG1 PC.^18^ Evidence from human B cells in vitro experiments suggested a delayed or impaired terminal differentiation of IgE PC compared to IgG1 PC ^67^. IgE PC from circulation,^15,16^ and from nasal polyps of allergic patients,^17^ were described as immature. Long-lived BM IgE PC were however described in mice after chronic antigen stimulation through the airways with house dust mite extract, ^33,41^ and in spleen and BM in OVA/alum immunized mice chronically challenged through the airways.^32^

Here, we demonstrated that long-lived IgE PC do form, they can be detected in mice several months after their formation, and they reside preferentially in the spleen and not the BM. Since new IgE PC form mainly in a restricted window of about a month after immunization, subsequent analysis of IgE PC detected mostly mature phenotypes that decreased in numbers overtime. The small number of long-lived IgE PC found after several months post immunization, confirmed to the mature phenotype of Ki-67^-^MHCII^low^CD93^+^CD98^high^ PC in spleen and BM.

Our studies of IgE PC biology described here demonstrate that IgE PC undergo maturation and terminal differentiation through the acquisition of an expression program that while sharing features common to other PC, displays distinct adaptations to cellular stress and survival. Using timestamping, we demonstrate the existence of long-lived IgE PC. Our findings have implications for the understanding of the mechanisms involved in persistent allergy and for the development of cell targeting therapies in allergy.

## RESOURCE AVAILABILITY

### Lead contact

Further information and requests for resources and reagents should be directed to and will be fulfilled by the Lead Contact, Maria A. Curotto de Lafaille.

## Material availability

All unique materials generated in this study are available from the Lead Contact with a completed Materials Transfer Agreement.

## Data and Code availability

The raw data for experiments performed, including flow cytometry files, scRNaseq data, are available upon request from the lead contact.

## ACKNOWLEDGMENTS

We thank Pedro Silva and Ankita Prakash for mouse colony management and general laboratory support; the Human Immuno Monitoring Center and the Genomics and Flow Cytometry Cores at ISMMS for technical support; Juan Lafaille and members of M.A.C.L. laboratory for critically reading the manuscript. Cartoons were created using Biorender. Work in the M.A.C.L. laboratory was supported by NIH grants R01AI130343, R01AI151707, and R01AI153708.

## AUTHOR CONTRIBUTIONS

M.C.G.M.W. and M.A.C.L. conceptualized the study and designed the experiments. M.C.G.M.W. carried out the experiments and performed analysis. E.C.A. helped with experiments. E.G.K. and L.X. performed the computational analysis of gene expression. K.B.H. performed BCR analysis. C.J.A. generated the *Prdm1^creERT^*^2^ mice. Y.G.C. helped with the AMINIS experiment. E.G.M.M. performed the initial experiments leading to the project. M.C.G.M.W. and M.A.C.L. wrote the manuscript. M.A.C.L. supervised the project. All authors read and approved the manuscript.

## DECLARATION OF INTERESTS

The authors declare no competing interests.

## SUPPLEMENTAL INFORMATION

Document S1. Figures S1-S8

Table S1. Excel file containing differentially express genes (DEG) of clusters, related to Figure 1.

Table S2. Excel file containing DEG from IgE PC and IgG1 PC maturation analysis, related to Figure 2.

## Notes

### Competing Interest Statement

The authors have declared no competing interest.

