## Supplemental document for "Long-lived IgE plasma cells that reside in the spleen contribute to the persistence of the IgE response"

**SUPPLEMENTARY FIGURES**

**Figure S1**

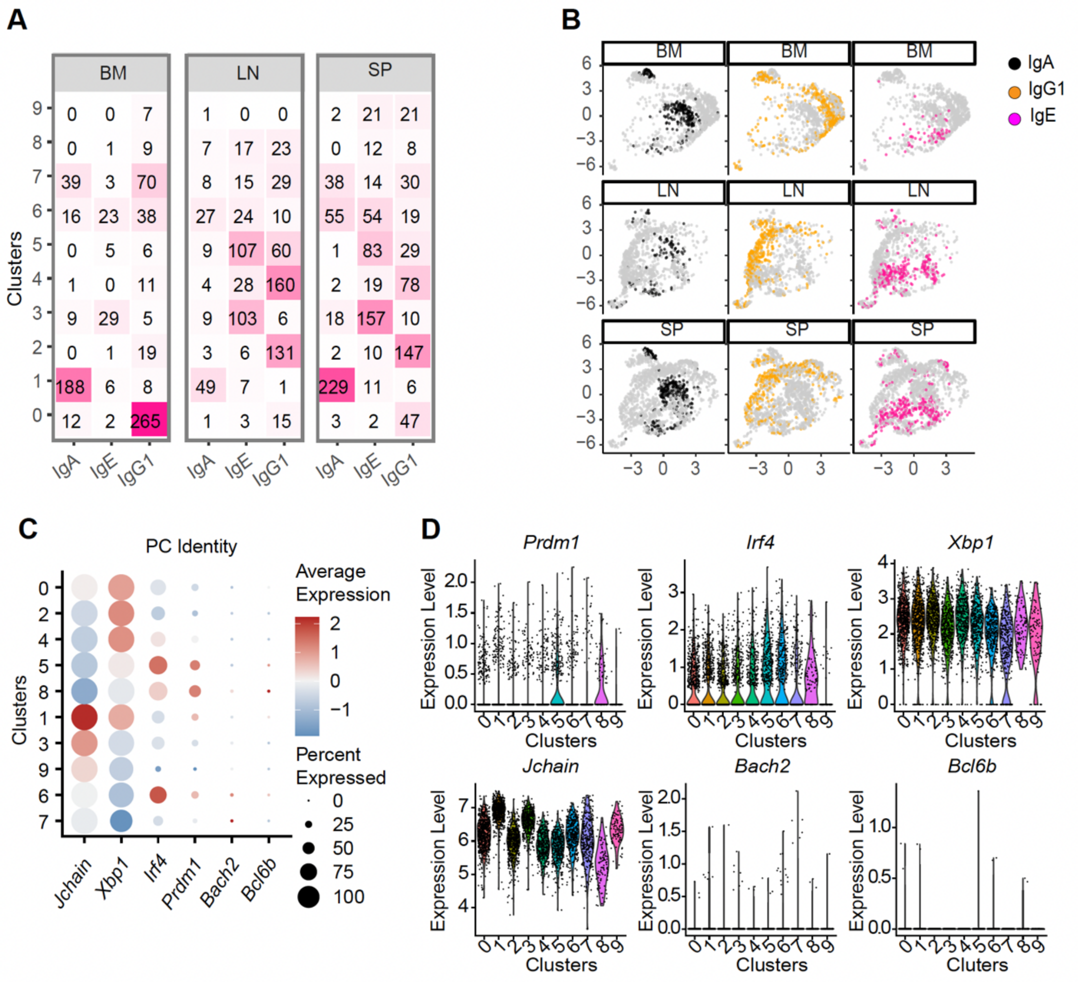


**Figure S1. Cell identity and tissue distribution of PC from immunized mice, analyzed by scRNAseq, related to Figure 1**

(A) Number of PC expressing IgA, IgE or IgG1 in each cluster and per tissue: bone marrow (BM), mesenteric lymph nodes (LN) and spleen (S).

(B) Projection of IgA, IgG1 and IgE PC from BM, LN and S on the UMAP of all PC.

(C-D) Gene expression of *Prdm1, Irf4, Xbp1* and *Jchain*, and of B cell genes *Bach2* and *Bcl6* in the cell clusters. (C) Dot plot graphics and (D) Violin plots.

### Figure S2

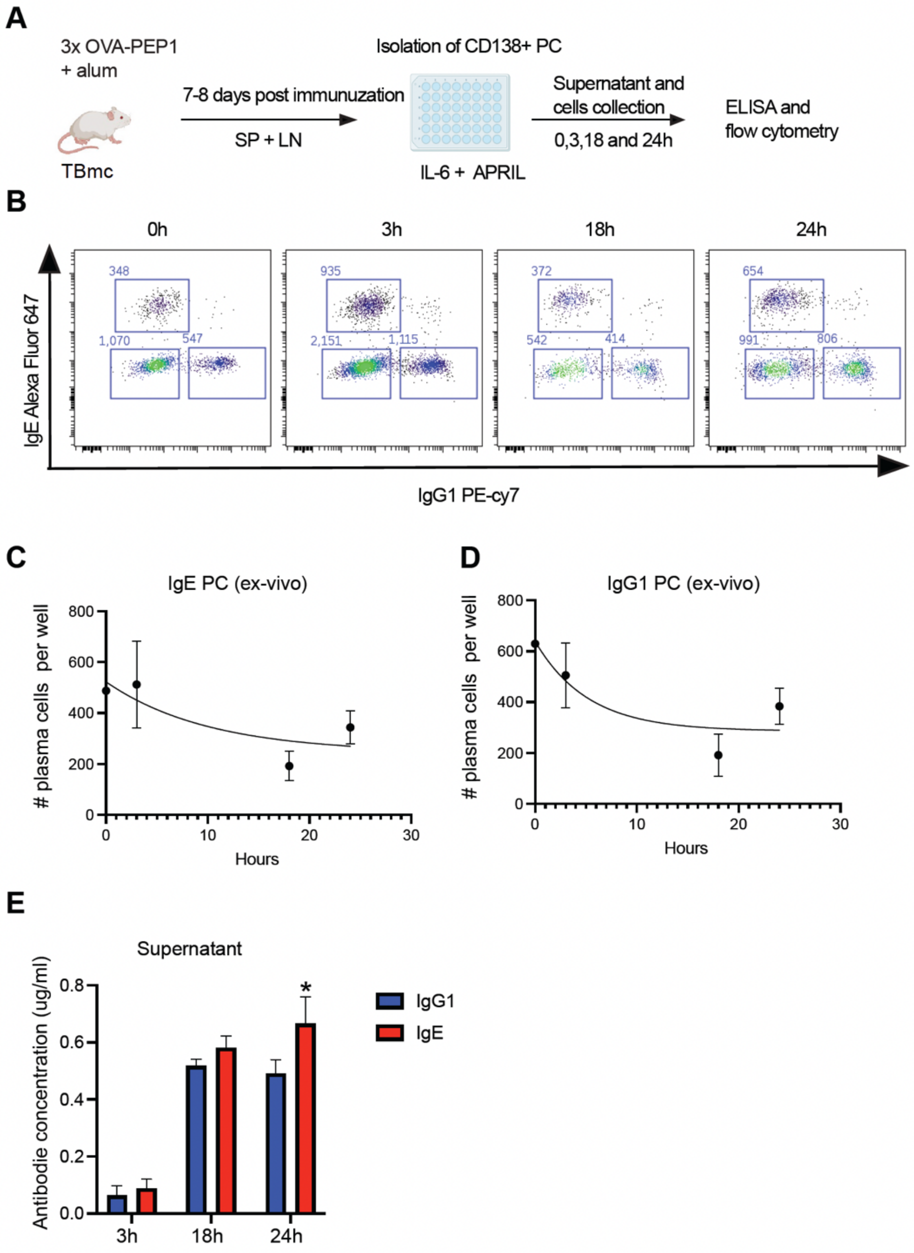


**Figure S2. Antibody secretion by IgE and IgG1 PC in vitro, related to Figure 2**

(A) Experimental approach. TBmc mice were immunized 3 times with OVA-PEP1 in alum. One week after the final immunization, PC were isolated from the spleen (SP) and mesenteric lymph nodes (LN) using anti-CD138 magnetic beads and cultured in RPMI medium supplemented with 10% FBS, IL-6, and APRIL. Supernatant and cell pellets were collected at 0, 3, 18, and 24 hours of culture for ELISA and flow cytometry analysis.

(B) Dot plots showing staining of IgE and IgG1 PC at 0, 3, 18, and 24 hours of culture.

(C) Quantification of IgE PC numbers per well over time.

(D) Quantification of IgG1 PC numbers per well over time.

(E) Concentrations of IgE and IgG1 antibodies in supernatants at 3, 18 and 24 hours of culture.

(C, D, E) PC were isolated from 8 mice, and three wells were analyzed per time point. Statistical analysis was performed using two-way ANOVA; *: indicates significant p-values ($\leq 0.05$). Shown mean ± SEM.

**Figure S3**

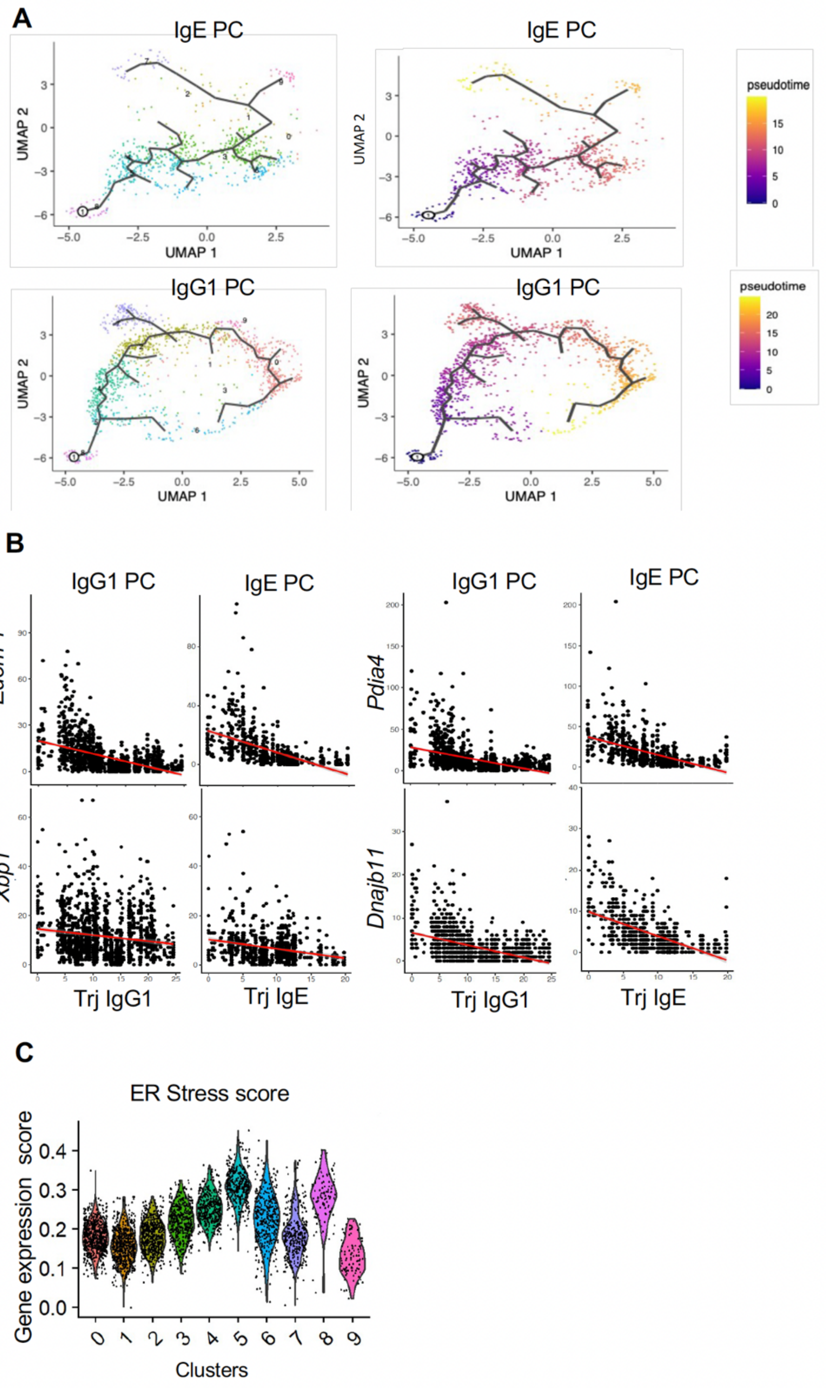


**Figure S3. Attenuation of ER stress with PC maturation correlates with high pseudotime values, related to Figure 2**

(A) UMAP representation colored by pseudotime, based on Monocle3 unsupervised graph and pseudotime analysis. The upper panel shows IgE plasma cells (PC), while the lower panel shows IgG1 plasma cells. The starting point was defined as the plasmablast cluster 8, and the lines indicate the predicted trajectory pathways.

(B) Correlation between gene expression and trajectory values in IgE and IgG1 PC.

(C) Violin plots showing gene signature scores for ER stress in the clusters.

**Figure S4**

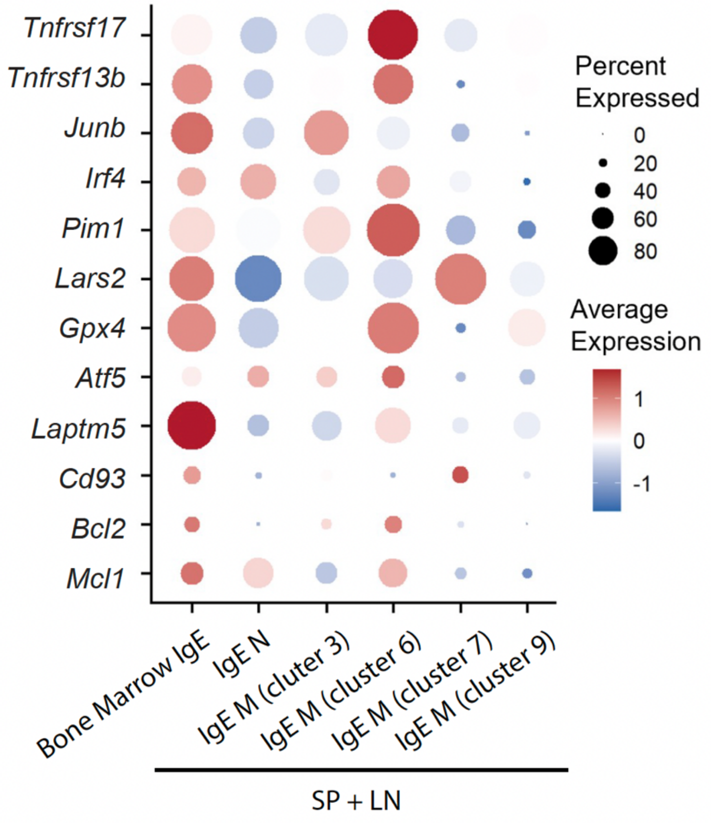


**Figure S4. High expression of survival genes in a subpopulation of mature IgE PC from the spleen, related to Figure 3**

Dor plots showing the expression of genes associated with PC survival in IgE PC from clusters containing bone marrow or spleen+LN IgE PC.

### Figure S5

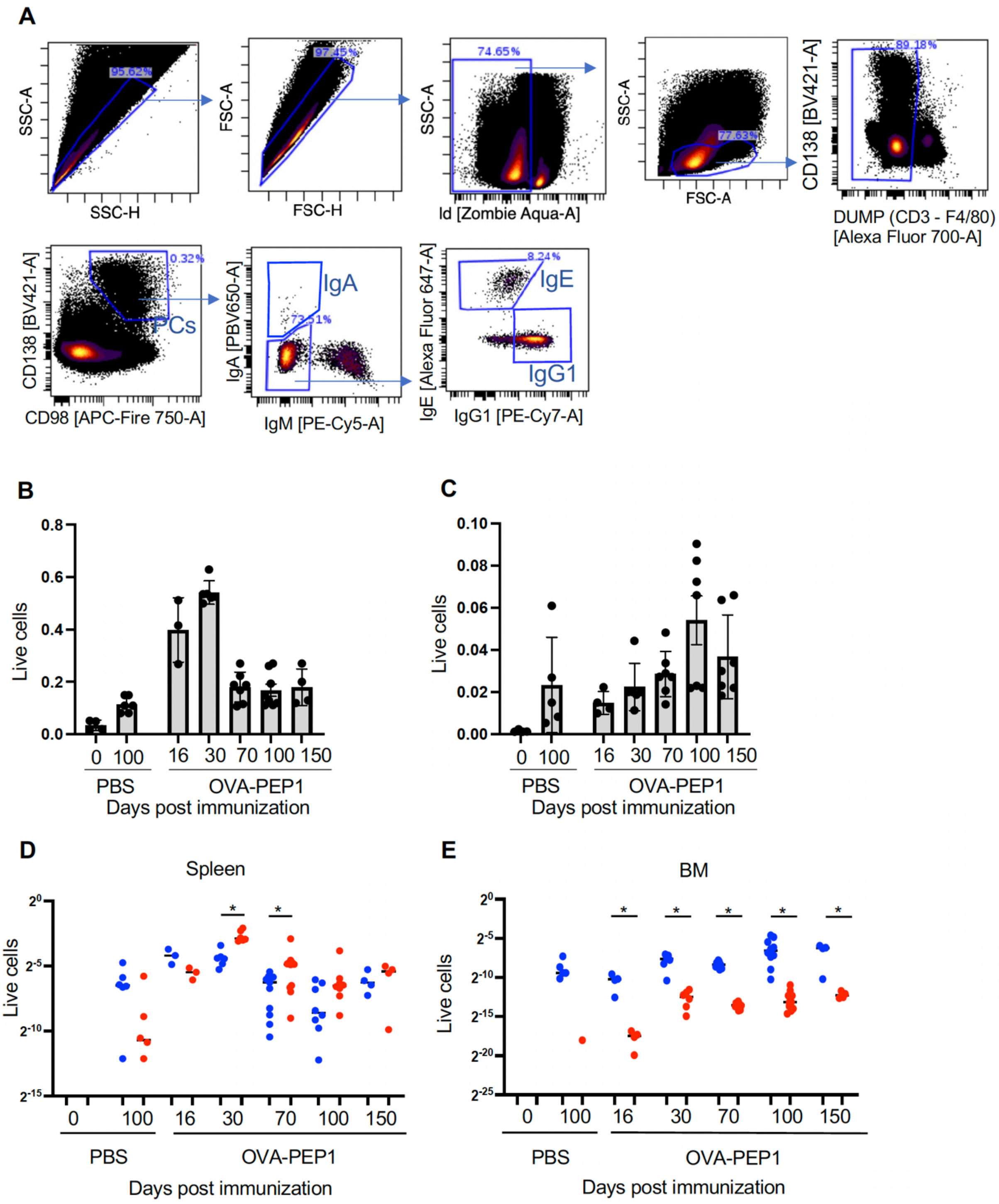


**Figure S5. Variations in PC numbers after a single immunization, related to Figure 4**

(A-E) TBmc mice were immunized with OVA-PEP1 in alum by intraperitoneal route. PC in spleen and bone marrow were analyzed at 16, 30, 70, 100 and 150 days after immunization and in control unimmunized mice (PBS treated).

(A) Gating strategy for PC.

(B) Frequencies of PC among live cells in spleen.

(C) Frequencies of PC among live cells in the bone marrow.

(D) Frequencies of IgE and IgG1 PC among live cells of the spleen.

(E) Frequencies of IgE and IgG1 PC among live cells of the bone marrow.

(B-E) Each dot represents a sample from an individual mouse. Statistical analysis comparing IgG1 and IgE PC at each time point was performed using an unpaired Student's *t*-test. Significant p-values ($\leq 0.05$) are indicated by *. Shown mean ± SEM.

**Figure S6**

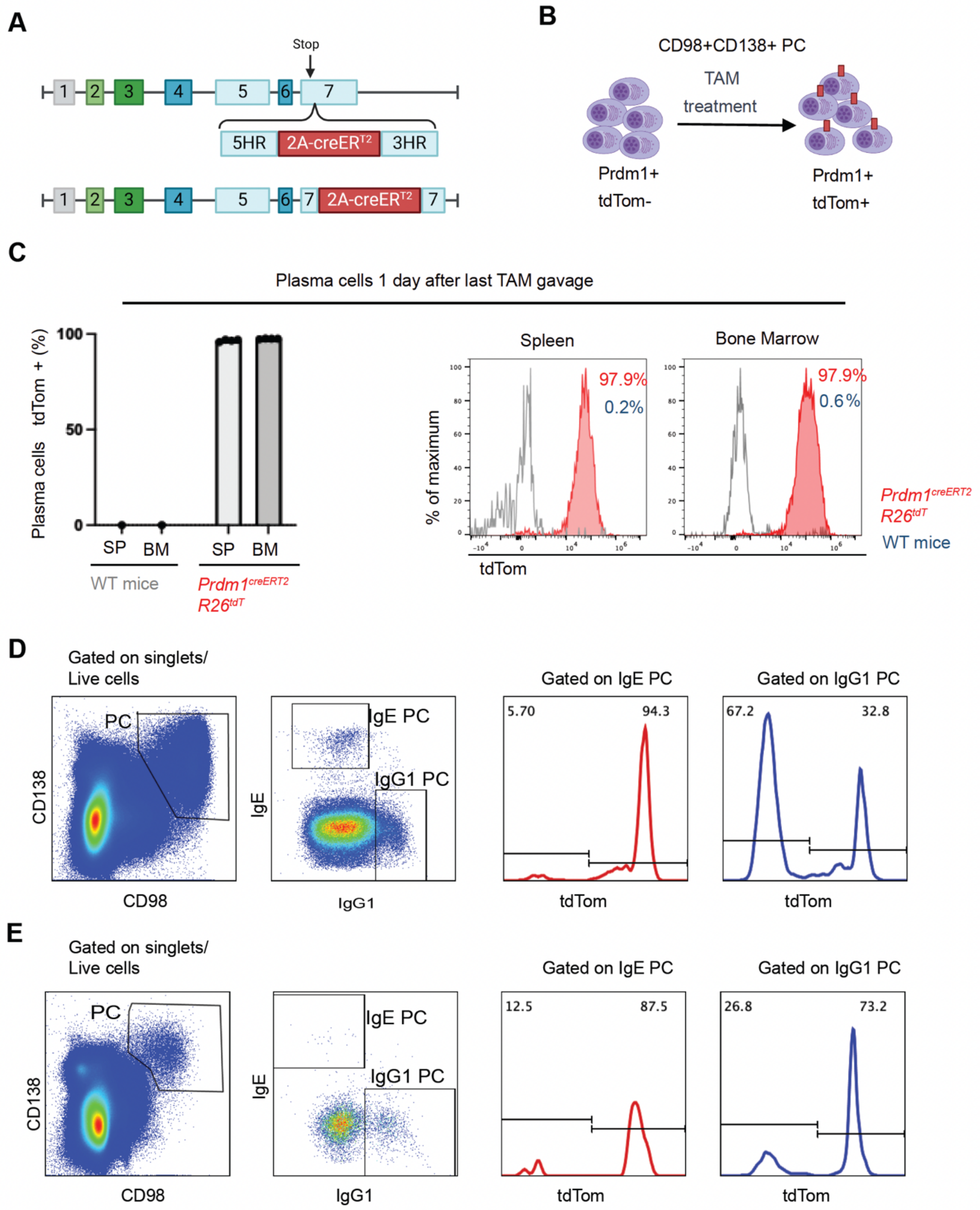


**Figure S6. Labeling of PC with tdTomato in *Prdm1*$^{CreERT2}$*R26*$^{tdT}$ mice, related to Figure 4**

(A) Generation of *Prdm1*$^{CreERT2}$*R26*$^{tdT}$ mice. *Prdm1*$^{CreERT2}$ mice were generated by inserting a *2A-creERT2* gene sequence downstream of the stop codon of the *Prdm1* gene using CRISP/Cas9 methodology. *Prdm1*$^{CreERT2}$ mice were then crossed with *Rosa26-stop-tdTomato* (*R26*$^{tdT}$) mice, in which a loxP-flanked transcriptional stop is located 5' of the *tdTomato* gene in the *Rosa26* locus.

(B) Upon tamoxifen treatment, most PC acquire tdTomato fluorescent expression.

(C) Wild type (WT) mice and $Prdm1^{CreERT2}R26^{tdT}$ mice were administered tamoxifen by gavage. One day later, the percentage of tdTomato+ PC among total PC in spleen and bone marrow were quantified. Most PC of spleen and bone marrow of $Prdm1^{CreERT2}R26^{tdT}$ express tdTomato, demonstrating effective timestamping of PC in these mice.

(D-E) $Prdm1^{CreERT2}$ $R26^{tdT}$ mice were infected subcutaneously with *Nippostrongylus brasiliensis* on days -30 and 0 and injected on days 5 and 6 with tamoxifen (TAM) to timestamp PC.

(D) Gating strategy to determine frequency of total and tdTomato+ PC in spleen.

(E) Gating strategy to determine frequency of total and tdTomato+ PC in bone marrow.

**Figure S7**

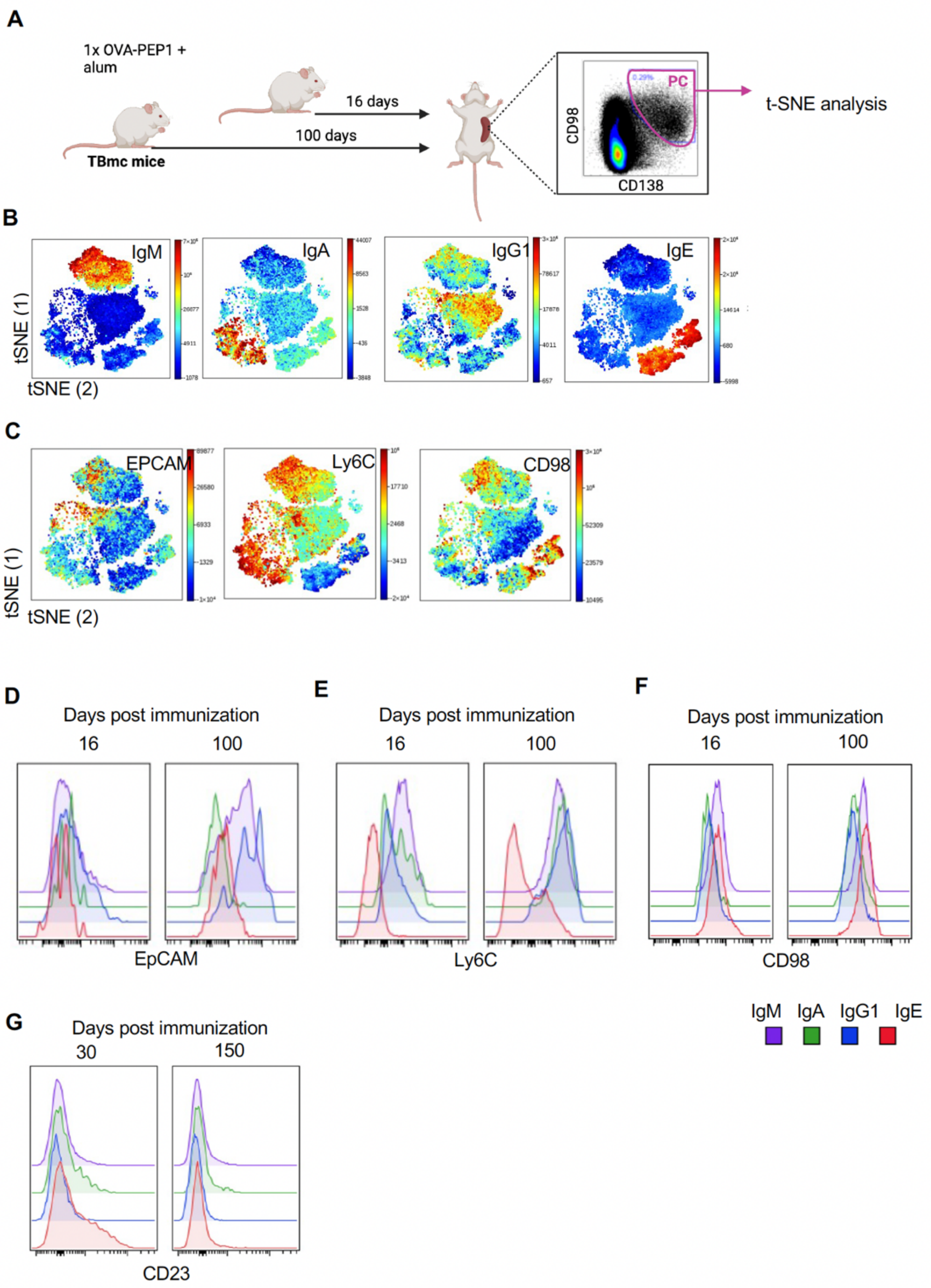


**Figure S7. Identification of maturation stages of IgE PC by spectral cytometry, related to Figure 5**

(A) Experimental approach. TBmc mice were immunized with OVA-PEP1 in alum 16 or 100 days before euthanasia. Total spleen cells were stained and analyzed using the Cytek Aurora spectral flow cytometer. opt-SNE analysis was performed on gated $CD138^{+}CD98^{+}$ total PC.

(B-C) t-SNE plots of concatenated spleen PC from 16 and 100 days post immunization.

(B) t-SNE plots showing expression of IgM, IgA, IgG1, and IgE.

(C) t-SNE plots showing expression of EpCAM, Ly6C, and CD98.

(D-G) Histograms displaying fluorescence expression of EpCAM (D), Ly6C (E), CD98 (F), and CD23 (G) in IgM, IgA, IgG1 and IgE cells at 16 and 100 days post immunization. Each histogram was obtained by concatenating data from 4 mice per timepoint.

**Figure S8**

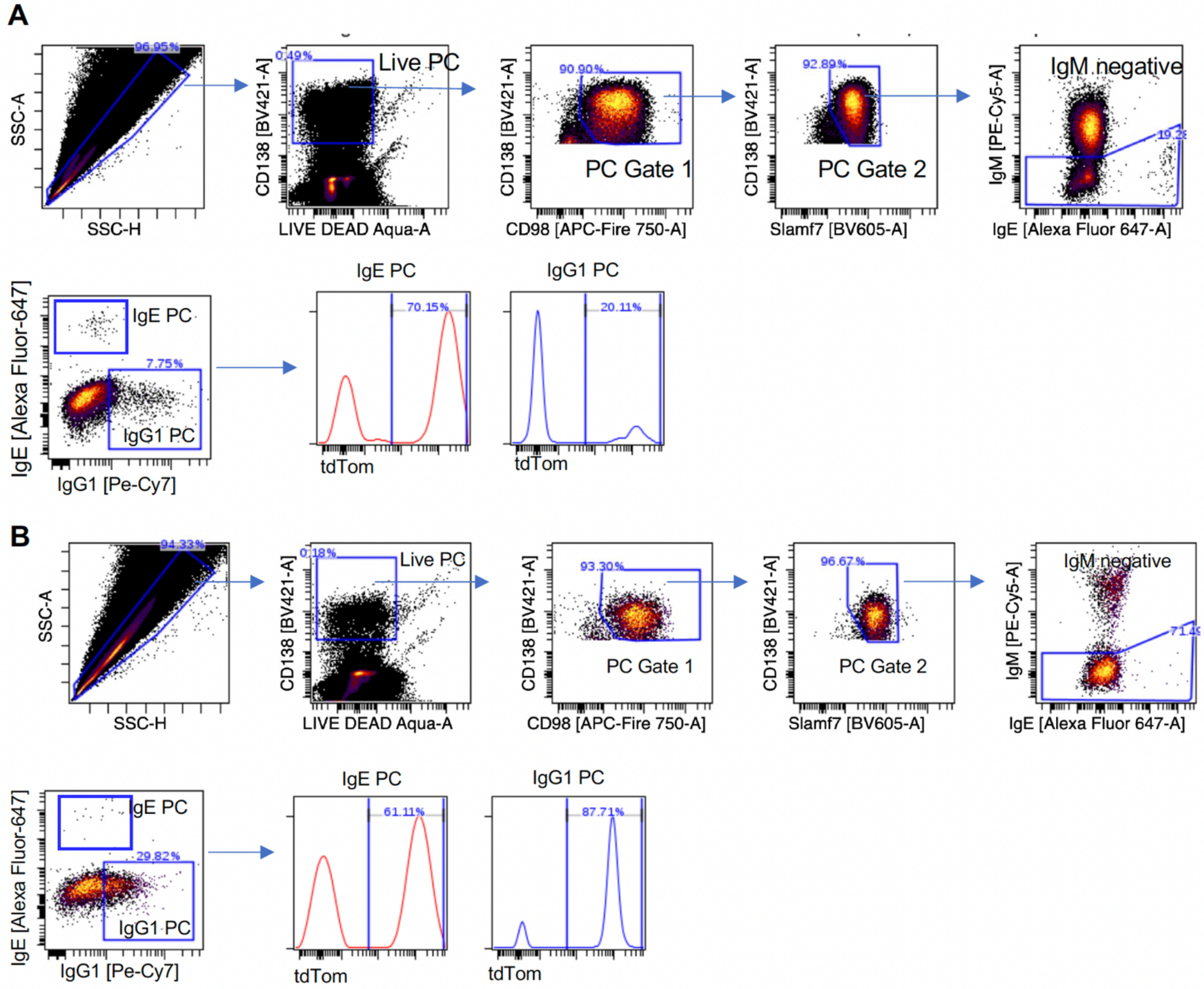


**Figure S8. Identification of long-lived PC by timestamping, related to Figure 7**

*Prdm1*$^{CreERT2}$ *R26*$^{tdT}$ mice were chronically exposed to intranasal *Alternaria alternata* extract for 10 weeks and were then administered tamoxifen (TAM) every other day during the following week. Mice were euthanized at 1 and 20 weeks post-tamoxifen. CD138 enriched PC were isolated from immunized mice. (A-B) Gating strategy of tdTomato+ IgE and IgG1 PC from spleen (A) and from bone marrow (B).
